# *marmite* defines a novel conserved neuropeptide family mediating nutritional homeostasis

**DOI:** 10.1101/2022.12.12.520095

**Authors:** AP Francisco, I Tastekin, AB Fernandes, G Ezra-Nevo, B Deplancke, AJ Oliveira-Maia, AM Gontijo, C Ribeiro

**Affiliations:** Champalimaud Foundation, Lisbon, Portugal; NOVA Medical School, Faculdade de Ciências Médicas, NMS, FCM, Universidade Nova de Lisboa, Lisbon, Portugal; Laboratory of Systems Biology and Genetics, Institute of Bioengineering, School of Life Sciences, Ecole Polytechnique Fédérale de Lausanne, Lausanne, Switzerland; iNOVA4Health, NOVA Medical School, Faculdade de Ciências Médicas, NMS, FCM, Universidade Nova de Lisboa, Lisbon, Portugal

## Abstract

Neuropeptides play a key role in regulating physiology and behavior, including feeding. While animals modify their food choices to respond to the lack of specific nutrients, the mechanisms mediating nutrient-specific appetites remain unclear. Here, we identified *marmite* (*mmt*), a previously uncharacterized *Drosophila melanogaster* gene encoding a secreted peptide that controls feeding decisions. We show that both *mmt* mutants and neuronal knockdown of *mmt* specifically increased the intake of proteinaceous food, whereas neuronal *mmt* overexpression reduced protein appetite. *mmt* expression is also higher in animals maintained on amino acid rich food, suggesting that *mmt* encodes a protein-specific satiety signal. Mmt is expressed in a small number of neurons in the adult nervous system, with a single pair of neurons modulating protein appetite. Finally, sequence and phylogenetic analysis showed that *mmt* is part of an ancient and conserved family of neuropeptides, including the poorly understood vertebrate *neuropeptides B* and *W* genes. Functional experiments showed that *mmt* and vertebrate *NPB* and *NPW* modulate food intake in both flies and mice. Therefore, we discovered an ancient family of neuropeptides involved in controlling feeding across phyla.

## Introduction

Neuropeptides and peptide hormones are important signaling molecules involved in almost all aspects of an animal’s life, from physiology to development and behavior (Andermann & Lowell, 2017; Nässel & Zandawala, 2019; Schoofs et al., 2017). They play vital roles in the dynamic adaptation of animals to multiple internal and environmental challenges and act at different temporal and spatial scales, ranging from cellular to systemic effects. These regulatory molecules are produced in neurons and neuroendocrine cells of the nervous system, enteroendocrine cells in the intestine or several peripheral sites. As such, they play an essential role in the capacity of animals to maintain homeostasis and maximize fitness. Given their essential and highly specialized role, peptide hormones are thought to be both ancient and fast evolving (Elphick et al., 2018). Together with their small size, it is however difficult to identify them using classic bioinformatics or experimental approaches, such as proteomics (DeLaney et al., 2018). Therefore, it is likely that new neuropeptides and peptide hormones remain to be discovered, including ones that are evolutionarily conserved across species.

A key aspect of homeostasis is maintaining a balanced intake of nutrients (Leopold & Perrimon, 2007; Münch et al., 2020; Simpson et al., 2015). Choices such as which food to eat and how much to ingest greatly impact the overall fitness of animals. Macronutrients such as proteins (amino acids), lipids and carbohydrates are essential for maintaining tissues and energy balance. Because they cannot be synthesized by the animal, essential amino acids (eAAs) are vital nutrients determining animal fitness, mainly through their effect on reproduction and aging (Leitão-Gonçalves et al., 2017; Piper et al., 2014). Despite their importance, overconsumption of AAs is known to reduce lifespan and negatively impact fitness (M. E. Levine et al., 2014; Piper et al., 2011; Solon-Biet et al., 2014). Therefore, the intake of this macronutrient must be tightly regulated and animals, including humans, have evolved mechanisms to modify their food choices to avoid the lack or overconsumption of AAs. Accordingly, in *Drosophila melanogaster* females, AA deprivation, mating, and the microbiome are potent modulators of protein intake (Ezra-Nevo et al., 2020; Henriques et al., 2020; Leitão-Gonçalves et al., 2017; Malita et al., 2022; Ribeiro & Dickson, 2010; Walker et al., 2015). Both nutrient and reproductive states have recently been shown to profoundly alter how taste information is processed in the fly brain leading to changes in protein appetite (Münch et al., 2022; Steck et al., 2018). Several neuropeptides and peptide hormones have been shown to regulate different aspects of the feeding process. Molecules signaling either satiety or hunger, reducing or promoting feeding, respectively, have been identified in several species, from flies to vertebrates (Andermann & Lowell, 2017; Lin et al., 2019). Neuropeptide Y in mammals and its orthologue Neuropeptide F in *Drosophila* are examples of two conserved peptides involved in promoting feeding (Brown et al., 1999; Horio & Liberles, 2021). However, signals regulating nutrient-specific appetites and how they act to control food choice remain less well understood.

In vertebrates, two neuropeptides that have been proposed to control feeding are the poorly studied neuropeptide B (NPB) and neuropeptide W (NPW), which share structural and functional similarities (Brezillon et al., 2003; Fujii et al., 2002; Shimomura et al., 2002; Takenoya et al., 2010; Tanaka et al., 2003). NPB and NPW are expressed in brain regions previously related to feeding, such as hypothalamic areas. Furthermore, intracerebroventricular administration of NPB or NPW in rodents has been shown to lead to changes in food intake (Aikawa et al., 2008; Baker et al., 2003; A. S. Levine et al., 2005; Mondal et al., 2003; Naso et al., 2014; Samson et al., 2004; Shimomura et al., 2002; Tanaka et al., 2003). Moreover, *NPB* knock-out mice have been reported to develop adult-onset obesity (Kelly et al., 2005). However, the precise function of NPB and NPW remains enigmatic as some studies observed an increase in food intake upon NPB or NPW injection (Baker et al., 2003; A. S. Levine et al., 2005; Samson et al., 2004; Shimomura et al., 2002), while others observed a reduction (Aikawa et al., 2008; Mondal et al., 2003; Naso et al., 2014; Tanaka et al., 2003). Therefore, while NPB/W signaling seems to play a role in controlling food intake and homeostasis, the mechanisms underlying their mode of action remain unclear. Likewise, the evolutionary origin and history of NPB/NPW remain unknown. Identifying evolutionary orthologues could open the door to a comparative mechanistic dissection of the function of this poorly understood peptide class and hence a deeper understanding of appetite regulation.

In this study, we identify *marmite*, a previously uncharacterized *Drosophila melanogaster* gene that encodes a predicted secreted peptide that controls protein appetite. Sequence and phylogenetic analysis show that this novel peptide is part of a conserved family of genes that likely originated early in the evolution of Bilateria and includes the vertebrate *neuropeptides B* and *W*. To characterize the behavioral relevance and anatomical distribution of *marmite* (*mmt*), we genome-engineered flies to generate animals either lacking *mmt* or harboring a *Gal4* sequence instead of the original coding region. *mmt* mutants and flies with neuronal knockdown of *mmt*, specifically increased their intake of proteinaceous food, while neuronal overexpression of *mmt* led to a decrease in protein appetite. Moreover, single essential AA deprivation decreases *mmt* expression, supporting the idea that *mmt* encodes a neuronal protein-specific satiety signal. This effect is corroborated by detailed anatomical and behavioral analysis, which showed that *mmt* is expressed in around 25 neurons in the brain, of which one pair is responsible for the protein appetite phenotype. Finally, we provided evidence that Marmite (Mmt) and NPB/NPW function is evolutionarily conserved: neuronal overexpression of human *NPB* or *NPW* genes in flies phenocopied the behavioral effect of *mmt* overexpression; and the overexpression of *mmt* in the mouse ventromedial hypothalamus (VMH) mimicked the overexpression of *NPB*. We therefore, identified a new AA-state controlled neuropeptide gene in *Drosophila* and showed that it encodes a satiety factor controlling protein intake via the action of one pair of neurons. Together with the vertebrate *NPB/NPW* genes, *mmt* defines an ancient neuropeptide family sharing a conserved function in controlling feeding.

## Results

### *CG14075* encodes a novel putative *Drosophila* neuropeptide

Despite the importance of neuropeptides in modulating behavior and physiology (Nässel & Zandawala, 2019), it remains difficult to predict these molecules and their corresponding genes. One of the challenges is their small size and poor sequence conservation, making it difficult to use existing proteomics and/or bioinformatics pipelines to identify all neuropeptides encoded by genomes. While the *Drosophila* genome is among the best-annotated genomes (Adams et al., 2000), many genes - mainly those with small open reading frames - encode products with unknown functions. One such gene is *CG14075*. It is predicted to en-code a product of 155 amino acids with no known or predicted domains (Gramates et al., 2022). Sequence analyses (see Materials & Methods) revealed a predicted signal peptide (SP) in the amino (N)-terminus and polybasic sites in the central part of the protein (Fig. 1a). Both are stereotypical features of neuropeptide precursors: the SP directs the peptide to the secretory pathway, while prohormone convertases typically recognize the identified basic residue motifs for proteolytic processing, a critical step in the generation of functional bioactive peptides (Pauls et al., 2014). These features strongly suggest that *CG14075* encodes an uncharacterized peptide which, given its expression in the nervous system (Li et al., 2022), could act as a novel neuropeptide regulating behavior.

**Figure 1.**
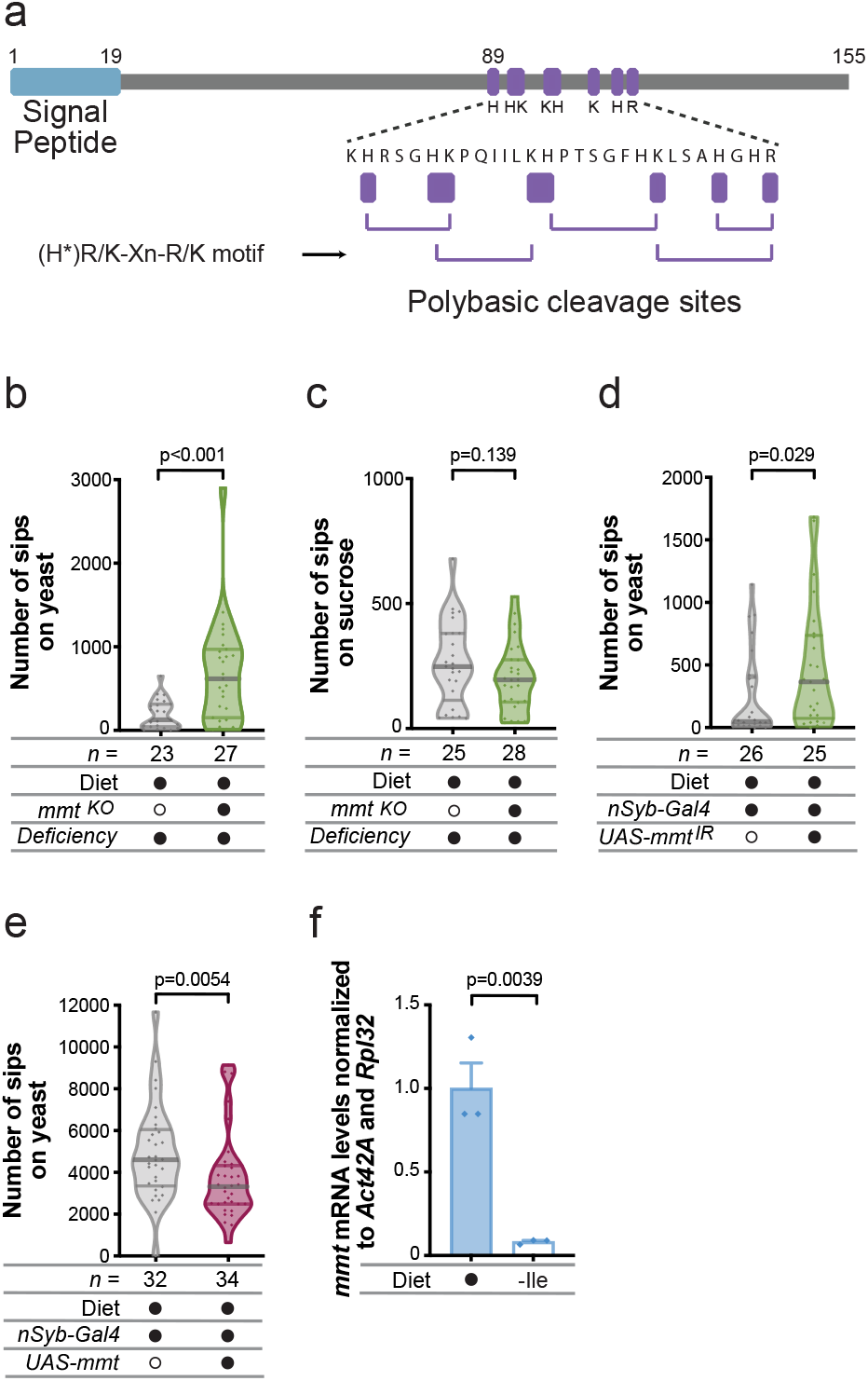
*mmt* is necessary and sufficient to alter protein appetite. **a)** Scheme depicting the 155 amino acid sequence of CG14075/Mmt with predicted motifs: signal peptide in the N-terminus and polybasic residues as potential targets for protein processing. The cleavage sites predicted by the motif (H*)R/K-Xn-R/K are shown, where X is any amino acid, n is 0, 2, 4, or 6 amino acids, and an H* denotes that histidine is accepted when n is 2, 4, or 6. The sequence and the size of the different motifs are represented to scale. The numbers on top indicate the AA position. **b)** Number of sips on yeast and **c)** sucrose of *mmt* mutant or control flies maintained on complete holidic medium. “Deficiency” denotes a genetic deficiency spanning the *mmt* gene. In this and the following figures, the number of sips was measured for each fly presented with a choice between feeding on yeast or sucrose, using the flyPAD system. The number of sips from each food source was measured separately. **d)** Number of sips on yeast of *mmt* pan-neuronal knockdown or control flies maintained on complete holidic medium. **e)** Number of sips on yeast of flies overexpressing *mmt* pan-neuronally or control flies maintained on holidic medium lacking Isoleucine (-Ile). **f)***mmt* mRNA levels measured from heads of flies maintained on complete holidic medium or holidic medium lacking Isoleucine (-Ile) as normalized using the expression levels of two control genes (*Actin 42A* and *RpL32*); columns represent the mean and the error bars the standard error of the mean. **b-e)** Below the graphs, filled black circles represent the complete holidic medium or the presence of a specific genetic element, whereas an empty black circle represents its absence. Violin plots show the frequency distribution of the data, with lines representing the median, upper and lower quartiles; points indicate the number of sips on yeast **(b, d-e)** or sucrose **(c)** for each fly. Significance was tested using the MannWhitney test. The number of flies used in each condition is shown in each panel. In **f)**, significance was tested using the unpaired *t*-test, n = 3.

### *mmt/CG14075* is necessary and sufficient to alter protein feeding

To address the putative functions of *CG14075*, we generated a knock-out allele using CRISPR-Cas9-based genome engineering (Suppl. Fig. 1, see also Materials & Methods). More specifically, we used two single guide RNAs (sgRNAs) to generate a double cut in the genome upstream and downstream of the *CG14075* coding sequence, leading to the replacement (and hence deletion) of the entire open reading frame of *CG14075* by an attP landing site via homologous recombination. We, therefore, generated a null allele of *CG14075* while allowing for later insertions of transgenes in the *CG14075* locus.

Given that neuropeptides play essential roles in controlling food intake (Andermann & Lowell, 2017; Lin et al., 2019), we decided to test for phenotypes in feeding behavior and food choice using the flyPAD technology, a capacitance-based setup for testing independent changes in protein and carbohydrate feeding (Itskov et al., 2014). Indeed, when given a choice to eat from a protein (yeast) or carbohydrate (sucrose) food source, fully fed *CG14075* mutant female flies showed a significant increase in protein feeding when compared to control animals (Fig. 1b), while sucrose feeding was not affected (Fig. 1c). We confirmed this phenotype using a video-tracking foraging setup allowing us to quantify the feeding behavior of flies in an arena containing protein and carbohydrate food spots (Corrales-Carvajal et al., 2016). Similarly to what we had observed using the flyPAD system, throughout the foraging assay, *CG14075* mutants showed a significantly longer total duration of micromovements (a proxy for feeding behavior) on yeast spots compared to controls (Suppl. Fig. 2a). Furthermore, we did not observe a phenotype for sucrose spots (Suppl. Fig. 2b). These results suggest that *CG14075* encodes a protein-specific satiety factor, whose loss leads to overconsumption of proteinaceous food. Given this phenotype, we named this gene *marmite (mmt)*, referring to the vital source of protein appreciated in some cultures.

**Figure 2.**
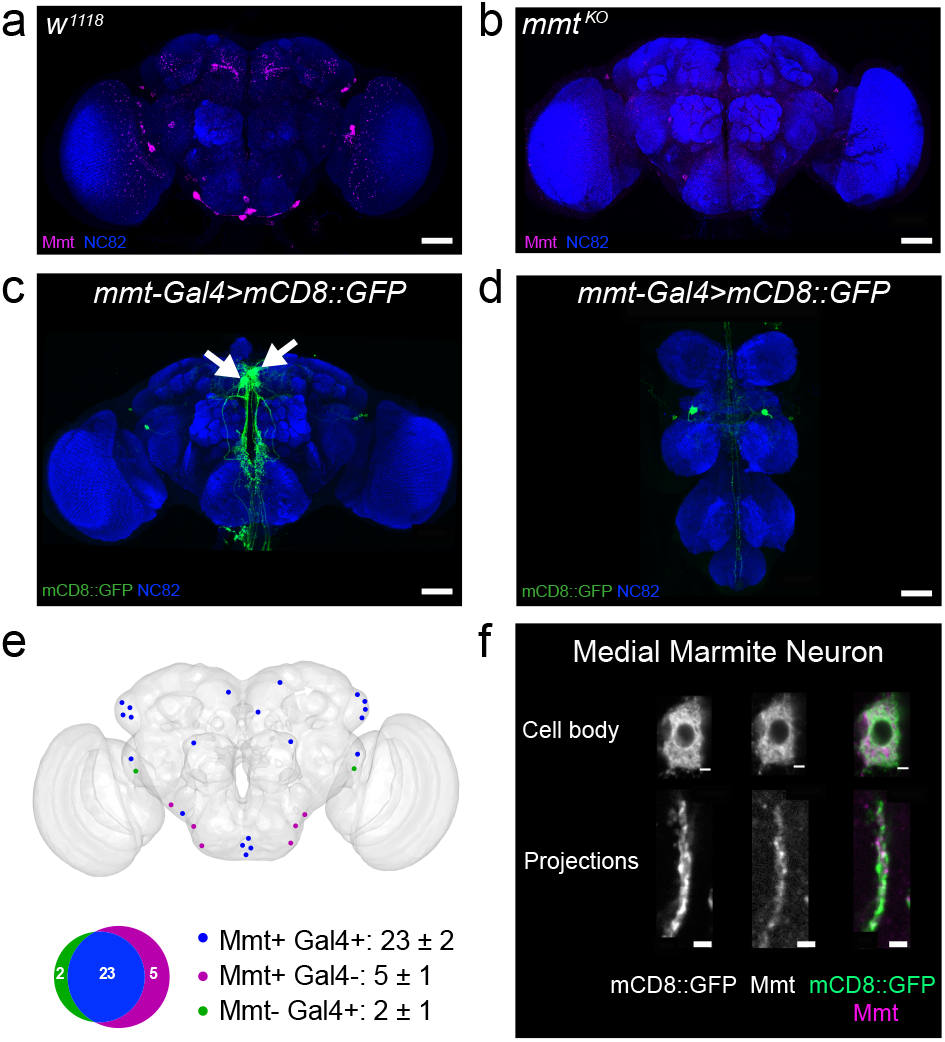
*mmt* is sparsely expressed in the nervous system. Mmt antibody staining of brains of (**a**) control and (**b**) *mmt* mutant flies in magenta, with nc82 neuropile staining in blue. (**c**) Expression pattern of *mmt*-*Gal4* in the brain and (**d**) VNC as visualized using *UAS-mCD8∷GFP* in green, with nc82 neuropile staining in blue. Arrows mark the Medial Marmite Neurons (MMNs). **e)** Schematic of a representative brain (see also Suppl. Fig 6) with the location of the soma of neurons showing labeling by both Mmt antibody and *mmt*-*Gal4* (Mmt + Gal4 +, blue), Mmt antibody only (Mmt + Gal4 -, magenta), or *mmt*-*Gal4* only (MmtGal4 +, green) as well as a Venn diagram depicting the overlap of these populations. Mean ± SEM for each combination is indicated, n=9. **f)** Single plane of confocal images of the cell body and projections of the MMNs showing mCD8∷GFP and Mmt labeling; scale bar corresponds to 2 μm. **a-d)** Brains are oriented with the dorsal side on top and the ventral side at the bottom. VNC is oriented with the anterior side on top and the posterior side at the bottom. Maximum projection images of confocal stacks; scale bars correspond to 50 μm.

Neuropeptides are peptides expressed in neurons that play crucial roles in regulating behavior and physiology. To test if *mmt* acts as a neuropeptide, we assessed if neuron-specific knockdown of *mmt* was sufficient to phenocopy the effects of the *mmt* knock-out on protein appetite. Indeed, knocking down *mmt* specifically in the nervous system led to a strong increase in the number of sips on yeast, strengthening the idea that it encodes a neuropeptide regulating protein appetite (Fig. 1d and Suppl. Fig. 3). If *mmt* encodes a neuropeptide mediating protein-specific satiety, overexpressing it should reduce protein appetite. This effect should be more prominent in females deprived of an eAA as these show a high protein appetite (Leitão-Gonçalves et al., 2017; Ribeiro & Dickson, 2010). Indeed, neuronal overexpression of *mmt* in mated females deprived of the eAA Isoleucine (Ile) led to a significant decrease in the number of sips on yeast when compared to control Ile-deprived females (Fig. 1e). These data lend further support to the idea that Mmt acts as a satiety signal dedicated to specifically reducing protein intake. Given this role, we would expect that its expression could be modulated by the AA state of the animal, which is a crucial signal controlling protein appetite (Henriques et al., 2020; Leitão-Gonçalves et al., 2017; Steck et al., 2018). We, therefore, compared mRNA expression levels of *mmt* in heads of flies kept on either a complete diet or a diet lacking Ile. In agreement with a role in linking AA state to protein satiety, Ile deprivation led to a decrease in *mmt* expression (Fig. 1f). These results strongly suggest that *mmt* encodes a neuropeptide whose expression is regulated by the animal’s AA state and that it enables the fly to alter protein appetite according to its nutritional needs.

**Figure 3.**
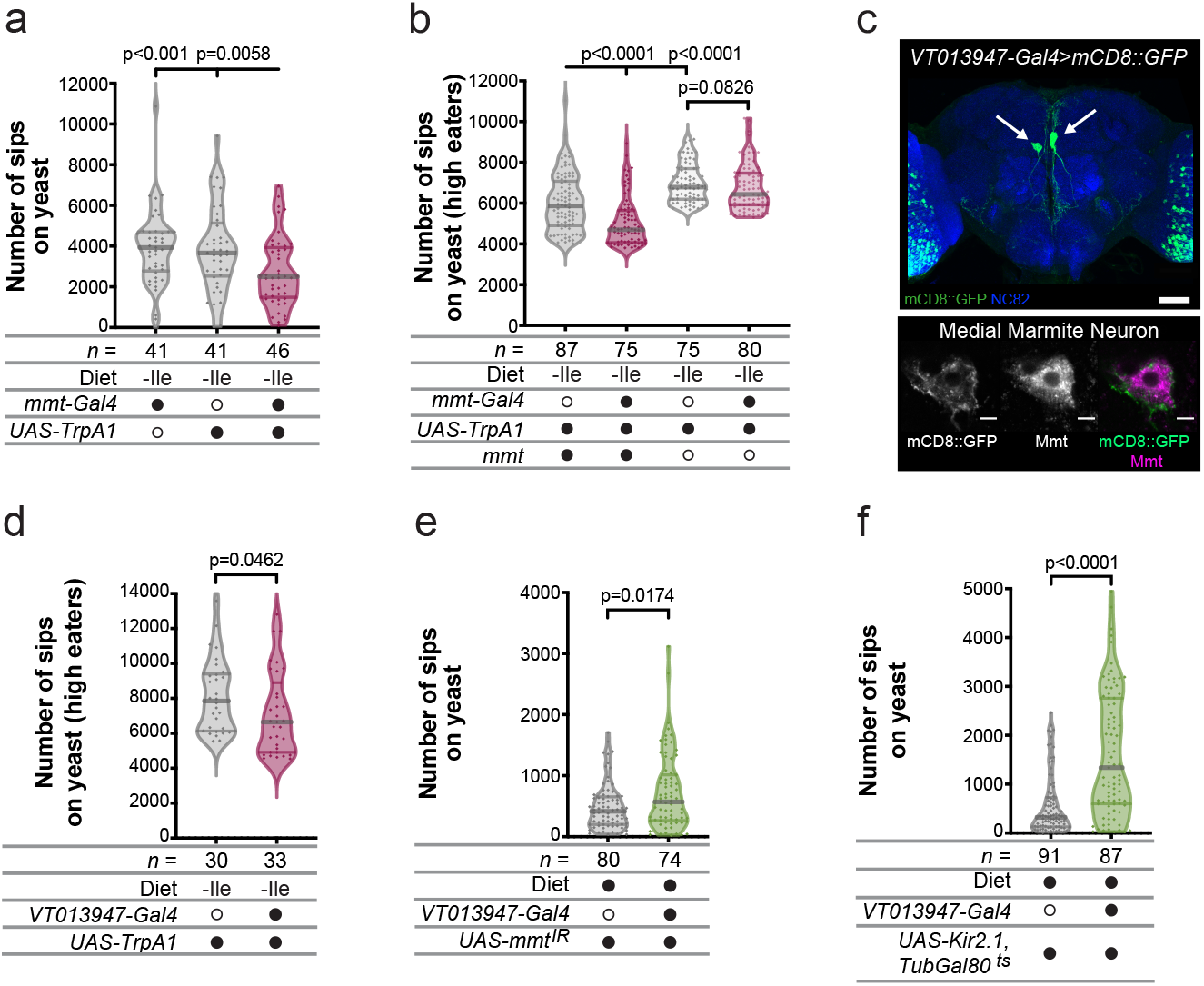
Silencing and activation of *mmt*-expressing neurons alters protein feeding. **a)** Number of sips on yeast of flies in which *mmt*-*Gal4* neurons were thermogenetically activated using TrpA1 and corresponding controls. Flies were maintained on holidic medium lacking Isoleucine (-Ile). **b)** Number of sips on yeast of flies in which *mmt*-*Gal4* neurons were thermogenetically activated using TrpA1 in a context of *mmt* presence or absence and corresponding controls. Flies were maintained on holidic medium lacking Isoleucine (-Ile). **c)** Top: Expression pattern of *VT013947*-*Gal4* in the brain visualized using *UAS-mCD8∷GFP* in green, with nc82 neuropile staining in blue; arrows highlight Medial Marmite Neurons; maximum projection of confocal images, scale bar corresponds to 50 μm. Bottom: single plane of one of the medial cell bodies showing mCD8∷GFP and Mmt labelling; scale bars correspond to 5 μm. Brain and soma are oriented with the dorsal side on top and the ventral side at the bottom. **d)** Number of sips on yeast of flies in which *VT013947-Gal4* neurons were thermogenetically activated using TrpA1 and corresponding control. Flies were maintained on holidic medium lacking Isoleucine (-Ile). **e)** Number of sips on yeast of flies in which *mmt* was knocked down in *VT013947-Gal4* and control flies. Flies were maintained on complete holidic medium. **f)** Number of sips on yeast of flies in which *VT013947-Gal4* neurons were acutely silenced using Kir2.1 and corresponding control. Flies were maintained on complete holidic medium. **a), b), d-f)** Below the graphs, filled black circles represent the complete holidic medium or presence of a specific genetic element, whereas an empty black circle represents its absence. Violin plots show the frequency distribution of the data, with lines representing the median, upper and lower quartiles; points indicate the number of sips on yeast for each fly. Significance was tested using the Mann-Whitney test. The number of flies used in each condition is shown in each panel.

### *mmt* is sparsely expressed in the nervous system

Neuropeptides are usually expressed by specific, specialized, and sparsely distributed neurosecretory neurons. To identify the neurons expressing the Mmt peptide, we first used a custom-generated polyclonal antibody to visualize the distribution of the peptide in the nervous system. Mmt was sparsely expressed in the central nervous system, labeling 28 cell bodies in the brain (Fig. 2a and 2e). Cell bodies could be found in the Subesophageal Zone (SEZ), the Posterior Ventrolateral Protocerebrum (PVP), the Lateral Horn (LH), and the Superior Medial Protocerebrum (SMP). Mmt was also expressed in 8 neurons in the Ventral Nerve Cord (VNC) (Suppl. Fig. 4a). Importantly, in *mmt* mutant animals, most of the staining was abolished, validating both the loss of function nature of the mutant as well as the high specificity of the antibody (Fig. 2b and Suppl. Fig. 4b).

**Figure 4.**
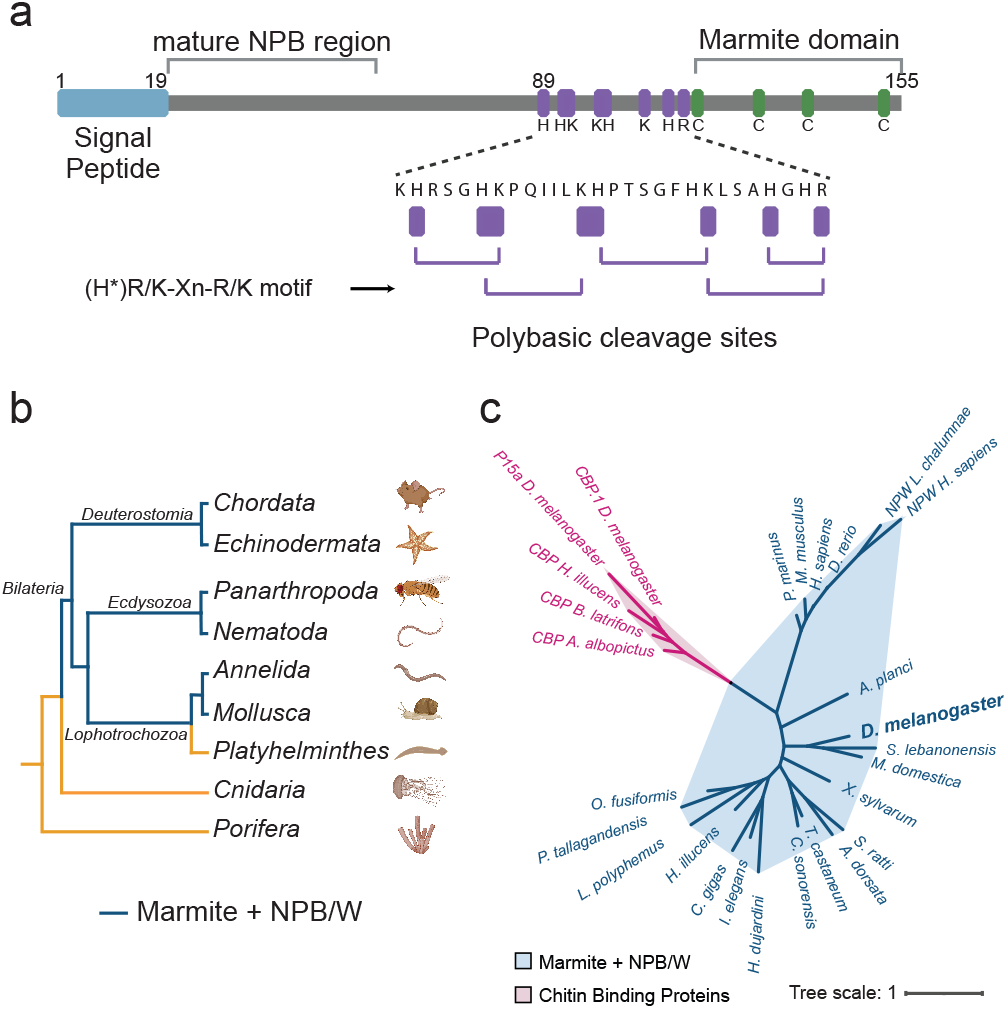
Mmt is part of a novel neuropeptide family which includes NPB and NPW. **a)** Scheme depicting the 155 amino acid sequence of the Mmt prepropeptide with predicted motifs: signal peptide in the N-terminus, region corresponding to the vertebrate mature NPB, polybasic residues as potential targets for protein processing and the four conserved cysteines in the C-terminus (Marmite domain). The cleavage sites predicted by the motif (H*)R/K-Xn-R/K are shown, where X is any amino acid, n is 0, 2, 4, or 6 amino acids, and an H* denotes that histidine is accepted when n is 2, 4, or 6. The size of the different motifs and the sequence are represented to scale. The numbers on top indicate the AA position. **b)** Cladogram with representative phyla in the animal kingdom highlighting the taxa where Mmt or NPB/W sequences can be found in blue and absent in orange. Created with Biorender.com. **c)** Phylogenetic tree showing the evolutionary relationship between AA sequences of Mmt-like, NPB/W proteins and members of the chitin binding protein family. Mmt-like and NPB/W sequences form a specific cluster in blue, separated from the CBP cluster in magenta.

Additionally, to gain genetic access to the neurons expressing *mmt*, we took advantage of the landing site inserted into the *mmt* locus in the null allele (Suppl. Fig. 1) and generated a knock-in allele with a *Gal4* inserted in the *mmt* locus (*mmt*-*Gal4*) (Suppl. Fig. 5). The expression of the *Gal4* should therefore mimic the endogenous expression of *mmt*. Accordingly, mCD8∷GFP driven by *mmt*-*Gal4* was expressed in neurons in the brain (Fig. 2c) and the VNC (Fig. 2d). To validate that the *mmt*-*Gal4* line faithfully recapitulates endogenous *mmt* expression, we visualized Mmt peptide expression in brains of flies in which *mmt*-*Gal4* drove mCD8∷GFP expression. Quantification of 9 brains revealed that the large majority of neurons (23±2) expressed both GFP and Mmt, with a small minority (5±1) expressing only the Mmt peptide and almost none being labeled by the Gal4 line and not the antibody (2±1) (Fig. 2e and Suppl. Fig. 6).

**Figure 5.**
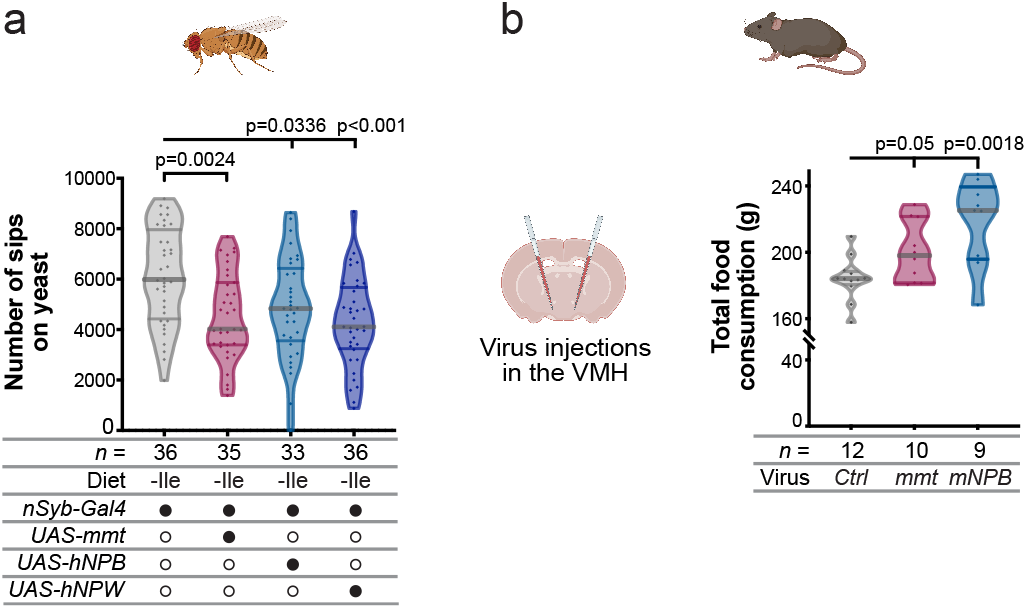
Mmt and NPB/W are functionally conserved. **a)** Number of sips on yeast of control flies and flies expressing *mmt*, the human *NPB* or *NPW* genes using a pan-neuronal driver. Flies were maintained on holidic medium lacking Isoleucine (-Ile). **b)** Left: scheme depicting the virus injection in the VMH of a mouse brain. Created with Biorender.com. Right: Total food consumption of mice injected with control virus or virus expressing either *Drosophila mmt* or mouse *NPB*. **a-b)** Below the graphs, filled black circles represent the presence of a specific genetic element, whereas an empty black circle represents its absence. Violin plots show the frequency distribution of the data, with lines representing the median, upper and lower quartiles; points indicate the number of sips on yeast for each fly **(a)** or the total food consumption for each mouse **(b)**. Significance was tested using the Mann-Whitney test (**a**) or unpaired *t-*test (**b**) with Bonferroni correction. The number of animals used in each condition is shown in each panel.

One of the most prominent labeled neurons was a pair of neurons with large somata in the medial part of the brain, which we named Medial Marmite Neurons (MMNs) (Fig. 2c and f). Upon close inspection, we could observe strong Mmt labeling in varicosity-like structures in the projections of these neurons (Fig. 2f). This staining pattern in the projections is compatible with Mmt being packed in vesicles, corroborating the idea that Mmt is a secreted neuropeptide.

Together, our data shows that Mmt is sparsely expressed in the central nervous system of the fly. In general, Mmt neurons have large cell bodies, which are a hallmark of neurosecretory cells. Given that we also observed Mmt labeling in varicosity-like structures within neuronal projections, our neuroanatomical analysis further supports the conclusion that *mmt* encodes a secreted neuropeptide.

### *mmt*-expressing neurons modulate protein feeding

Given the function of *mmt*, we hypothesized that changing the activity of neurons expressing the peptide should also lead to specific changes in feeding. We, therefore, took advantage of *mmt*-*Gal4* to activate these peptidergic neurons using the temperature-sensitive cation channel TrpA1 (Hamada et al., 2008). Since we expected this would reduce protein feeding, we decided to do this manipulation in Ile-deprived animals.

Consistent with previous results using *mmt* overexpression, activation of *mmt*-expressing neurons led to a decrease in the number of sips on yeast when compared to the controls (Fig. 3a). Given our results, it is likely that the neuronal activation effect on feeding is due to an increase in Mmt release in response to neuronal activation. To show that the *mmt* gene product mediates this effect, we decided to compare the effect of the neuronal activation experiment in an *mmt* expressing (while still being hemizygous as *mmt*-*Gal4* is an *mmt* null allele) and an *mmt*-null background. Importantly, *mmt* null mutants showed an increase in yeast feeding when compared to hemizygous controls, confirming the reproducibility of the *mmt* phenotype in a different genetic background and nutritional condition (Fig. 3b and Suppl. Fig. 7). This effect was even clearer when focusing on the top half of the distribution (high eaters) (Fig. 3b). This fits the rationale that flies with a low level of feeding would be less sensitive to the manipulation as they would already be subjected to a satiation signal. Importantly, whereas activation of *mmt*-*Gal4* in a control condition led to a significant decrease in yeast intake, the same activation in an *mmt* mutant background had no significant effect (Fig. 3b and Suppl. Fig. 7). These data show that the reduction in protein feeding observed upon activating *mmt* neurons is dependent on Mmt.

We have shown that *mmt*-*Gal4* labels 25 neurons in the brain (Fig. 2c and e). For multiple neuropeptides and neuromodulators, careful neurogenetic dissection has shown that it is possible to attribute specific behavioral functions to subsets of neurons (Liu et al., 2017; Waddell, 2013). We, therefore, set out to test if we could identify a subset of *mmt* neurons mediating the yeast satiety effect. We identified a driver line from the Vienna Tile (VT) collection (VT013947) (Tirian & Dickson, 2017) whose expression pattern comprises a pair of neurons that resembles the Medial Marmite Neurons (Fig. 3c and 2c). Acute thermogenetic activation using *VT013947-Gal4* reduced yeast feeding (Fig. 3d and Suppl. Fig. 8a). As observed earlier, the activation effect is stronger in animals with a higher level of yeast feeding (high eaters). Co-staining experiments with Mmt antibody showed that the medial neurons labeled by *VT013947-Gal4* are indeed Mmt positive and that no other neurons labeled by this *Gal4* line express Mmt (Fig. 3c and Suppl. Fig 8b). This finding strongly suggests that the medial neurons labeled by this *Gal4* line are the Medial Marmite Neurons. To further validate that the medial neurons labeled by *VT013947-Gal4* control yeast feeding via Mmt, we knocked down *mmt* expression using RNAi driven by this *Gal4* line and tested if this recapitulates the increase in yeast feeding observed upon pan-neuronal *mmt* knockdown (Fig. 1d). Indeed, loss of *mmt* function specifically in these neurons was sufficient to increase yeast appetite (Fig. 3e), consistent with the idea that Mmt suppresses yeast feeding via the Medial Marmite Neurons.

Additionally, acute silencing of the neuronal population labeled by *VT013947-Gal4* using Kir2.1 (Baines et al., 2001), an inward rectifying potassium channel, led to a significant increase in yeast appetite (Fig. 3f). This result mimics the effect of the *mmt* loss of function experiments and can be easily explained by a possible block of Mmt release due to the hyperpolarization of the neurons. This neuronal silencing effect is further supported by the significant increase in the total duration of yeast micromovements observed using the video-tracking setup (Suppl. Fig. 8c).

Altogether, our results show that the activity of *mmt*-expressing neurons controls protein feeding in an *mmt*-dependent manner and that a single pair of Mmt-positive neurons is mainly responsible for this effect.

### Mmt and vertebrate neuropeptide B/W define a new conserved neuropeptide family

Homeostatic regulation is a crucial feature of all living organisms. As such, many peptides, including neuropeptides, are conserved across evolution (Brown et al., 1999; Melcher et al., 2006). However, the entire repertoire of conserved neuropeptides is yet to be discovered as these are notoriously known to be highly divergent at the sequence level (Rajan & Perrimon, 2012). To understand the molecular evolutionary history of Mmt and gain further insight into possible critical residues, we first aligned it with Mmt-like proteins from other *Drosophila* and closely related insect species. Not surprisingly, we observed relatively poor amino-acid conservation across the protein sequence, with the notable exception of a conserved motif corresponding to four cysteine residues in the carboxy (C)-terminus of the prepropeptide (Fig. 4a and Suppl. Fig.9). Using this motif, which we named the Marmite domain, and filtering hits for those predicted to also encode a signal peptide and have a similar length (see Materials & Methods), we found a group of proteins in several vertebrate species corresponding to the peptide precursor (prepropeptide) of neuropeptide B (NPB), a known vertebrate neuropeptide (Brezillon et al., 2003; Fujii et al., 2002; Tanaka et al., 2003) (Suppl. Fig. 9). In vertebrates, this family of neuropeptides includes the *neuropeptide W* (*NPW*) gene (Shimomura et al., 2002; Tanaka et al., 2003), which encodes a prepropeptide with a very similar amino (N)-terminal peptide sequence to NPB. Our sequence analysis suggests that *NPW* has likely duplicated from an *NPB*-like ancestral gene. Intriguingly, NPW seems to have exchanged the conserved C-terminal region, containing the conserved cysteine residues, for a different C-terminal part (Suppl. Fig. 10).

Further sequence- and motif-based searches identified similar sequences in other invertebrates, including ecdysozoans (Nematoda and Panarthropoda), lophotrochozoans (Mollusca and Annelida), and deuterostomes (e.g., in Echinodermata) (Fig. 4b and Suppl. Fig. 9). Hence, *mmt*-like neuropeptide genes are found in all major bilaterian clades (Ecdysozoa, Lophotrochozoa, and Deuterostomia). Even though we did not find examples of Mmt-like sequences in some major clades (such as Platyhelminthes), the most parsimonious interpretation of these results is that an *mmt*-like gene must have been present in the last common ancestor of all bilaterians.

To rule out that the identified relationship between Mmt and NPB neuropeptides is spurious and/or both peptide families are part of an even larger family, we performed further relaxed sequence searches for other proteins with any resemblance to the identified Mmt/NPB C-terminal motif, retaining those hits that satisfied the criteria of being a putatively secreted protein in the hundred-AA-length range. This search identified insect members of a Chitin Binding Protein (CBP) family, which comprises chitinases and peritrophic matrix proteins (Shen & Jacobs-Lorena, 1999). The CBP domain of these proteins belongs to the CBM14 family of carbohydrate-binding domains, also known as peritrophin-A domain in insects (Chang & Stergiopoulos, 2015). However, two major pieces of evidence indicate that Mmt/NPB are part of a distinct family of proteins: first, CBP proteins are characterized by a 6-cysteine motif instead of the 4-cysteine motif found in Mmt/NPB, and a few highly conserved aromatic residues (notably at position-3 before cysteine 3, and position 6 after cysteine 4), none of which are present in Mmt/NPB precursors. Second, whereas a minority of predicted CBP proteins have 4 cysteines (e.g., CG14300 in *Drosophila melanogaster*), their sequence otherwise aligns well with the other CBPs (hence their detectability as CBPs in PFAM domain searches), which is not the case of Mmt/NPB precursors. Consistent with this, phylogenetic analysis showed that Mmt-like sequences and the vertebrate NPB and NPW form a separate cluster from CBP proteins, including those containing only 4-cysteines, supporting our finding that Mmt, NPB and NPW are part of a new, ancient, and conserved neuropeptide family (Fig. 4c). Thus, we propose to call this new family the Marmite Neuropeptide Family.

### Mmt and neuropeptide B/W are functionally conserved

Our phylogenetic analysis suggests that Mmt and NPB/W are part of the same conserved family of neuropeptides. Intriguingly, NPB and NPW have been implicated in feeding regulation in vertebrates (Aikawa et al., 2008; Baker et al., 2003; A. S. Levine et al., 2005; Mondal et al., 2003; Naso et al., 2014; Samson et al., 2004; Shimomura et al., 2002; Tanaka et al., 2003). This raises the possibility that the Mmt/NPB neuropeptide family has evolved to regulate feeding across species and that their conservation covers the functional level, extending beyond pure sequence conservation. Therefore, we decided first to test whether the human *NPB* and *NPW* gene products could alter feeding in *Drosophila*. To test for functional conservation of the encoded neuropeptides, we focused on phenocopying the decrease in yeast feeding observed upon neuronal *mmt* overexpression (Fig. 1e). Strikingly, pan-neuronal overexpression of either the human *NPB* or *NPW* led to a clear decrease in yeast intake, which mimicked the phenotype observed upon *mmt* overexpression (Fig. 5a). These results strongly suggest that the *mmt*, *NPB*, and *NPW* gene products are functionally conserved, as they can exert the same effect on feeding in flies. Furthermore, given that the NPW prepropeptides lack the Marmite domain (the cysteines in their C-terminal domain are not conserved) (Suppl. Fig. 10) suggests that the N-terminal part of the prepropeptide, just downstream of the SP, exerts the observed behavioral effect. Importantly, it is the N-terminal part that has been shown to give rise to the neuropeptides exerting the feeding effect in rodents, lending further support to our hypothesis (Aikawa et al., 2008; Baker et al., 2003; A. S. Levine et al., 2005; Mondal et al., 2003; Naso et al., 2014; Samson et al., 2004; Shimomura et al., 2002; Tanaka et al., 2003).

To further strengthen the argument for functional conservation, we decided to test if overexpression of the *Drosophila mmt* gene can induce changes in food intake in rodents. Intracerebroventricular administration of either NPB or NPW peptides has been shown to alter food intake in rodents (Aikawa et al., 2008; Baker et al., 2003; A. S. Levine et al., 2005; Mondal et al., 2003; Naso et al., 2014; Samson et al., 2004; Shimomura et al., 2002; Tanaka et al., 2003). We therefore, decided to use a targeted gain-of-function strategy to test if Mmt can alter feeding in mice. To do so, we engineered AAV viruses to neuronally express either the mouse *NPB* or the *Drosophila mmt* gene. For each group, we injected one of these viruses or a corresponding control bilaterally in the Ventromedial Hypothalamic nucleus (VMH), as this brain area has been shown to express NPB in mice (Fujii et al., 2002) (Suppl. Fig 11). Injected mice were kept in single cages and their food consumption was monitored for 8 weeks using a diet containing 14% protein. Consistent with what we observed in flies, overexpression of both *mmt* and *NPB* led to changes in feeding compared to controls (Fig. 5b). Animals overexpressing either of these neuropeptides’ genes showed a significant increase in cumulative food intake over 8 weeks when compared to controls. These data, therefore, suggest that the *Drosophila mmt* gene encodes a neuropeptide that can similarly alter food intake as NPB, supporting the idea that the function of the ancient Marmite Neuropeptide Family is conserved across evolution.

## Discussion

Neuropeptides are ancient regulatory molecules that play important roles in adapting biological processes to the animal’s internal and external world. While multiple conserved peptide families have been discovered, we are likely to still miss many important regulators of animal behavior and physiology. In this study, we discovered that *marmite*, a previously uncharacterized neuronal gene in *Drosophila*, encodes a neuropeptide that controls protein appetite. While Marmite is expressed in about 28 neurons in the central fly brain, we showed that one pair of neurons, the MMNs, is required and sufficient to mediate the protein satiety effect of Mmt. Furthermore, phylogenetic analysis showed that with the vertebrate neuropeptides B and W, Mmt defines a new family of ancient neuropeptides whose origin can be traced back to the beginning of bilaterians. Accordingly, functional studies showed that the biological activity of members of the Marmite Neuropeptide Family is conserved, as both the vertebrate and fly genes can alter feeding behavior in both flies and mice.

### Mmt acts as a protein-specific satiety signal

The nutritional state of an animal is in constant change. A variety of neuronal and peripheral signals induced by the lack (“hunger” signals) or sufficiency (“satiety” signals) of nutrients have been identified in animals, including *Drosophila*, and most of these signals correspond to neuropeptides and peptide hormones (Andermann & Lowell, 2017; Itskov & Ribeiro, 2013; Lin et al., 2019). Hugin, for example, a proposed fly homolog of the mammalian Neuromedin U, acts as a satiety signal, where high *hugin* expression in larvae is correlated to low food intake and reduced food-seeking behavior (Melcher & Pankratz, 2005). Other examples of satiety factors, such as FIT, MIP, DSK and Upd1, have also been described to limit food intake (Beshel et al., 2017; Carvalho-Santos et al., 2020; Min et al., 2016; Söderberg et al., 2012; Sun et al., 2017). While most studies focus on general satiety states, few nutrient-specific satiety signals are known (Münch et al., 2020). Our data suggest that Mmt functions as a protein-specific satiety signal in *Drosophila*. *mmt* mutants overconsume protein-rich food (yeast), while having no apparent phenotype for sucrose consumption, thus suggesting that the absence of *mmt* has a specific effect on protein but not carbohydrate appetite. This is supported by our observation that neuronal overexpression of *mmt* reduced yeast feeding, even in the context of eAA deprivation when flies have a strong drive to eat yeast. A third observation supporting this idea is that *mmt* expression is higher in fully-fed flies when compared to flies deprived of Ile. These data strongly support the idea that Mmt transforms the AA state of the fly into a neuronal signal which is then used to control protein intake. The discovery of *mmt* provides an opportunity to study important questions about feeding and protein homeostasis regulation. These include how internal AA availability is transformed into changes in *mmt* expression, what is the mature form of the processed Mmt peptide altering protein intake, and what is the receptor and concomitantly the target cells through which Mmt changes food choice. Additionally, animals have several satiety and hunger signals, some of which are nutrient-specific and some likely to control bulk food intake independently of the macronutrient content of the food. Once we have identified all the main players controlling nutrient homeostasis, a crucial but challenging task for the field will be to understand how Mmt signaling intersects with other neuronal and peripheral signals to fine-tune nutrient homeostasis across a wide range of states and temporal scales.

### *mmt*-expressing neurons also play a role in protein appetite

Most *Drosophila* neuropeptides are produced in a small subset of neurons or neuroendocrine cells. Peptidergic neurons often have stereotypically large cell bodies and extended axon ramifications (Nässel & Zandawala, 2019). Visualization of the peptide distribution using a custom-made Mmt antibody and membrane-targeted GFP expression driven by *mmt*-*Gal4* showed that the peptide is produced in a few neurons with large cell bodies and vesicle-like structures in their projections. These features further support that *mmt* encodes a secreted neuropeptide. Similar to what has been observed with other neuropeptidergic neurons, the behavioral effect of genetically manipulating the peptide gene can be partially mimicked by manipulating neuronal activity: thermogenetic activation of *mmt*-expressing neurons led to a reduction in protein feeding while inhibiting their activity led to the opposite effect. Importantly, this effect was abolished in an *mmt* mutant background. This shows that while these neurons might express other neuropeptides and neuromodulators, their effect on protein appetite is mediated by Mmt. Furthermore, these data suggest that the action of Mmt is regulated at multiple levels, including the regulation of *mmt* expression by the AA state of the animal, as well as the regulation of neuronal activity and neuropeptide secretion. This also explains why the effect of neuronal activation and inhibition were sometimes subtler than those observed when manipulating the gene. To avoid ceiling and floor effects, these experiments were done in conditions where the level of *mmt* expression was opposite to the desired effect, likely counteracting the impact of our neuronal manipulations.

Intriguingly, while around 28 neurons produce Mmt in the central brain, our experiments suggest that one pair of neurons is required and sufficient for mediating the protein appetite effect of the neuropeptide. These experiments suggest a specialization among *mmt* neurons, with the MMNs being important for mediating the protein appetite effect of Mmt. Similar findings have been made for other neuropeptidergic and neuromodulatory neurons, where anatomically distinct neurons seem to mediate specific effects (Liu et al., 2017; Waddell, 2013). This adds a further layer of regulation, allowing for the modulation of the effects of this neuropeptide, highlighting the complexity but also the sophistication of state-specific control of neuronal function and behavior by neuropeptides. The identification of MMNs as important mediators of protein-specific satiety will allow a focused dissection and identification of downstream circuits controlling protein appetite. These results also suggest that the other *mmt*-expressing neurons and, therefore, also Mmt, might exert further regulatory roles beyond controlling protein appetite. Careful neurogenetic, behavioral, and physiological analyses should allow for the identification of these roles in future studies.

### Mmt and vertebrate neuropeptide B/W define a new conserved neuropeptide family

Neuropeptides are signaling molecules that emerged early in animal evolution (Elphick et al., 2018; Nässel & Zandawala, 2019; Schoofs et al., 2017). Thus, it is not surprising that some neuropeptides have maintained similar sequences and functions, and that orthologues are found among different taxa. However, due to their small size and the conservation of only a few critical residues in most cases, establishing evolutionary relationships based on sequence queries has been challenging. Mmt and Mmt-like gene products in related insect species share a fully conserved motif of four cysteines in the C-terminal part of the protein (Marmite domain). This domain, together with the size range of the proteins and the presence of a signal peptide, led us to identify a group of vertebrate gene products with similar features, which corresponded to the NPB prepropeptide. Once processed, this previously characterized neuropeptide is very similar to NPW (Chottova Dvorakova, 2018). However, the prepropeptide of NPW, does not have the Marmite Domain. This could suggest that the precursor of NPW originated from a duplication of an NPB-like ancestral and its C-terminus diverged after that. Hence, even though the NPW prepropeptides have lost the Marmite domain, they share a common ancestor with Mmt and NPB prepropeptides. Moreover, we found Mmt/NPB-like sequences in other non-vertebrate taxa, strongly suggesting that the Mmt neuropeptide family is ancient and was already present in the common ancestor of all bilaterians.

Almost all members of the Marmite neuropeptide family have the Marmite domain. In general, this domain is by far the most strongly conserved portion of all members of the Marmite neuropeptide family (except NPW). Considering all of this evidence, we conclude that the Marmite domain is unique and most likely originates from evolutionarily-related genes with a separate evolutionary history from CB-Pencoding genes, at least since the evolution of Bilateria. This raises the question of what could be the function of this domain. Prepropeptides often encode multiple peptides with different biological activities (Jékely et al., 2018). In other cases, domains within prepropeptides play important roles in processing or chaperoning the peptide, which harbors the receptor binding and hence signaling activity (Vejrazkova et al., 2020). Currently, all functional studies in rodents have focused on the peptide encoded by the N-terminal region of NPB and NPW, leaving us with no information about the function of the conserved C-terminal region. Identifying the function of the Marmite domain will be an exciting avenue for future research.

### Mmt and neuropeptide B/W are functionally conserved

Our data on the functional characterization of the *mmt* gene and *mmt*-expressing neurons demonstrated the role of this peptide and the corresponding neurons in regulating food intake, specifically the appetite for protein-rich food. NPB and NPW neuropeptides have previously been linked to feeding and body weight regulation (Aikawa et al., 2008; Baker et al., 2003; Kelly et al., 2005; A. S. Levine et al., 2005; Mondal et al., 2003; Naso et al., 2014; Samson et al., 2004; Shimomura et al., 2002; Tanaka et al., 2003). We, therefore, tested whether the conservation between these vertebrate neuropeptides and Mmt could extend to the functional level. Indeed, in *Drosophila*, the neuronal overexpression of the human forms of both *NPB* and *NPW* led to a reduction in yeast intake, similar to when *mmt* was overexpressed. Likewise, in mice, the overexpression of the mouse *NPB* and *Drosophila mmt* genes in the brain led to increased feeding. These data suggest that similar to what has been observed for other peptides, Marmite Neuropeptide Family members can exert similar functions across vast evolution spans, even when bioinformatically, the conservation can only be pin-pointed to a few residues.

Which part of the product encoded by Marmite neuropeptide family genes produces the observed behavioral effect? In vertebrates, the N-terminal part of the prepropeptides encoded by *NPB* and *NPW* gives rise to NPB and NPW, respectively. Importantly, all rodent studies (including those studying feeding) focused exclusively on studying the function of these N-terminal-derived peptides (Aikawa et al., 2008; Baker et al., 2003; Mondal et al., 2003; Samson et al., 2004; Shimomura et al., 2002; Tanaka et al., 2003). The prepropeptide encoded by the *mmt* gene has several potential cleavage sites in the middle part of the sequence, just before the four-cysteine motif. At this stage, we do not know whether and how this precursor is cleaved and processed and what is the mature form of the bioactive peptide. However, given the results with the overexpression of the vertebrate peptides and the fact that in NPW, the Marmite domain is not conserved, our data suggest that for Mmt, similarly to vertebrates, it is the N-terminal part of the prepropeptide that is likely to produce a biologically active peptide which binds to a receptor to modify feeding.

While NPB and NPW have been repeatedly shown to alter feeding in rodents, how they do so and their exact effect remains unclear. Both anorexic (Aikawa et al., 2008; Mondal et al., 2003; Naso et al., 2014; Tanaka et al., 2003) and orexigenic (Baker et al., 2003; A. S. Levine et al., 2005; Samson et al., 2004; Shimomura et al., 2002) effects have for example been observed. While we observed a similar feeding phenotype upon *NPB* and *mmt* overexpression in the VMH, the fact that we observed a gain of feeding in mice and a reduction in feeding in flies might be interpreted as contradictory. However, it is important to note that this could be due to the complexities of NPB action or differences in the physiological role of NPB signaling in vertebrates and invertebrates. As indicated by the seemingly contradictory results in rodents (Samson et al., 2004; Tanaka et al., 2003), the mode of action of NPB might be complex. It could exert different effects depending on its target site in the brain (both the peptide and the receptor have a complex expression pattern), the exact internal state of the animal, and/or the nutrient content of the food options presented to the animal. In this sense, it is intriguing that we observe that *mmt* has a protein-specific effect in flies. If this is the case for vertebrates, then the animal’s exact protein, reproductive and microbiome states, and the protein content of the food might be important variables to be considered when interpreting specific effects of manipulating NPB. In this sense it is intriguing that gastric *NPW* levels in mice seem to be mainly responsive to the protein content of the food (Li et al., 2014). Nevertheless, the fact that both NPB and Mmt elicit the same effect in mice suggests that their biological activity is related. Our results indicate that a thorough understanding of how NPB and NPW act in the vertebrate brain will require a careful molecular and circuit dissection in the context of precisely controlled conditions, with specific attention to the protein content of the tested food. The ability to design parallel experiments in invertebrates and vertebrates should expedite such a functional understanding, be it at the level of inspiration or by using the strength of different organisms when deciding where to perform which experiment. Especially the ability for high throughput experiments in flies provides a substantial advantage for testing an increasing number of structure-function analyses.

### *Drosophila* as a discovery system for identifying novel regulators of homeostasis

*Drosophila melanogaster* was the second animal to have its genome fully sequenced. Much has been learned since then, and, one might be tempted to think that together with the extensive genetic screens performed on flies, we know the entire repertoire of crucial gene products regulating different traits. We hope that our discovery of *mmt* and the Marmite neuropeptide family is an inspiration for further digging in the gold mine that is the genome and biology of this organism. This is especially the case for molecules that help the animal manage deviations in homeostasis. It is likely that many fascinating and important molecular mechanisms will only reveal their existence and function when the animal is specifically perturbed or when using careful phenotypic analysis, as is required when studying behavior. From many existing regulatory systems controlling brain function, neuropeptides remain among the most difficult to probe. Their small size makes them difficult to discover, their multilayered regulatory mechanisms challenging to manipulate, and their mode of function difficult to parse. It is clear that we still lack tools and generalizable strategies to dissect neuropeptide function efficiently in animals. Nevertheless, their importance in regulating animal behavior and physiology makes them key players in understanding organismal function in health and disease. Advancing our understanding of these molecules should therefore be a top priority. Multi-phyla approaches have been vital in advancing our understanding of molecular processes from development to brain function. The advent of sophisticated molecular, neuronal, and genetic manipulation methods and the ability to use these in a vast repertoire of organisms make such comparative multi-phyla approaches feasible and more relevant than ever. Identifying new peptides such as Mmt, and dissecting their evolutionary relationship and mode of function is likely to be a fruitful, impactful, and fascinating research avenue for many years to come.

## ACKNOWLEDGMENTS

We thank Célia Baltazar, Ana Paula Elias and Inês Vicente for technical assistance at all stages of the project; Dennis Goldschmidt for analysis of the video-tracking data; Carolina Quadrado and Tatiana Saraiva for help with rodent behavioral measurements and image processing; Moita Lab (Ricardo Silva Neto), Vasconcelos Lab (Márcia Aranha), E. Chiappe, and the Jean-Paul Vicent Lab for providing fly stocks and reagents; the Champalimaud Fly and Rodent Platforms, the Molecular and Transgenic Tools, and the Glass Wash and Media Preparation Platforms for technical support throughout the project. We thank Joaquim Alves da Silva, Samuel J. Walker, Zita Carvalho-Santos and the whole Behaviour and Metabolism laboratory for helpful discussions and comments on the manuscript. Lines obtained from the Bloomington Drosophila Stock Center (NIH P40OD018537) and the VDRC were used in this study. We thank Ricardo Henriques for creating the LATEX template. APF was supported by a doctoral fellowship PD/BD/114277/2016 from the Portuguese Foundation for Science and Technology (FCT). IT was supported by Marie Skłodowska-Curie Actions postdoctoral fellowship MSCA-IF-EF-ST 867459. ABF was supported by a postdoctoral contract funded by FCT (SFRH/BPD/88097/2012; DL57/2016). GE-N was supported by a Marie Skłodowska-Curie Widening Postdoctoral Fellowship (101038098). BD was supported by Personalized Health and Related Technologies (PHRT) Grants (#2017/502 & #2019/719). AJO-M and ABF were supported by national funds through the FCT under the project PTDC/SAU-NUT/3507/2021 and by the European Research Council (ERC) under the European Union’s Horizon 2020 research and innovation program (grant agreement No 950357). AMG was supported by the FCT and UNL (Universidade Nova de Lisboa) under the CEEC Institutional program. Research in the AMG lab was supported by FCT (PTDC/BIA-BID/31071/2017; PTDC/MED-NEU/30753/2017; EXPL/BIA-BID/1524/2021; EXPL/BIA-COM/1296/2021) and by iNOVA4Health – UIDB/04462/2020 and UIDP/04462/2020 and by the Associated Laboratory LS4FUTURE (LA/P/0087/2020) financed by the FCT/MCTES (Portugal). The project leading to these results has received funding from FCT (FCT-PTDC/BIA-MOL/30081/2017) to CR.

Research at the Centre for the Unknown is supported by the Champalimaud Foundation and by Portuguese national funds through FCT - Fundação para a Ciência e a Tecnologia - in the context of the project UIDB/04443/2020, the research infrastructure CONGENTO, co-financed by Lisboa Regional Operational Programme (Lisboa2020), under the PORTUGAL 2020 Partnership Agreement, through the European Regional Development Fund (ERDF) and FCT under the project LISBOA-01-0145-FEDER-022170, and the PPBI - LISBOA-01-0145-FEDER-022122.

## Materials and Methods

### *Drosophila* rearing, media, and dietary treatments

Flies were reared on yeast-based medium (YBM) (per liter of water: 8 g agar [NZYTech, PT], 80 g barley malt syrup [Próvida, PT], 22 g sugar beet syrup [Grafschafter, DE], 80 g corn flour [Próvida, PT], 10 g soya flour [A. Centazi, PT], 18 g instant yeast [Saf-instant, Lesaffre], 8 ml propionic acid [Argos], and 12 ml nipagin [Tegospet, Dutscher, UK] [15% in 96% ethanol] supplemented with instant yeast granules on the surface [Saf-instant, Lesaffre]). To ensure a homogenous density of offspring among experiments, fly cultures were always set with 6 females and 3 males per vial and left to lay eggs for 3 d. Flies were reared in YBM until adulthood.

Holidic media (HM) were prepared as described previously (Leitão-Gonçalves et al., 2017) using the exome match formulation with food preservatives (Piper et al., 2017). For Holidic medium-Isoleucine (-Ile), Isoleucine was not added to the diet, keeping the concentration of all other nutrients constant. For all experiments using the HM, the following dietary treatment protocol was used to ensure a fully-fed state: groups of 1±5-d old flies (16 females) were collected into fresh YBM-filled vials, 5 *Canton-S* males were added to the vials to ensure females were mated, and transferred to fresh YBM after 48 h (or 24 h in flies kept at 22°C). After 24 h, flies were transferred to different HM for 3 days or 5 days (if flies are kept at 22°C for temperature-sensitive experiments) and immediately tested in the indicated assay. All experiments were performed with female flies. Flies were reared and maintained at 25°C in climate-controlled chambers at 70% relative humidity in a 12-h light-dark cycle (Aralab, Fito-Clima 60000EH). Behavioral testing was performed at 25°C or 30°C (for experiments using *UAS-TrpA1*). Polypropylene fly vials (VWR, #734±2261) were used.

### *Drosophila* stocks and genetics

*VT013947-Gal4* (VDRC #205650) and *CG14075* RNAi (VDRC #14328) were obtained from the VDRC stock center, and the deficiency spanning *CG14075* (*w^1118^; Df(3L)BSC220/TM6C, Sb^1^*, *cu^1^* - BDSC #9697) was obtained from BSDC. Other stocks and the genotypes of flies used in this study are listed in Table 1.

### Generation of *mmt* knock-out and *Gal4* knock-in lines

To generate both *mmt* knock-out line and the *Gal4* driver line, we adapted protocols from two-step CRISPR- and RMCE-based strategies for genome editing in *Drosophila* (Port & Bullock, 2016; Zhang et al., 2014). To generate the knockout line (Suppl. Fig 1), we first cloned 2 single guide RNAs (sgRNAs) into a pCFD5 plasmid (Addgene #73914): sgRNA1 atcgtccggcagccagcgttcgg (189bp upstream of the start codon) and sgRNA2 tcaatcaggccgcaagtgaa (26bp downstream of the stop codon). We used the pTV3 (gift from Jean-Paul Vicent Lab) as a vector for the donor DNA, including an attP site (that will be inserted into the locus for later use) and *mCherry* as a selective marker under the 3xP3 promoter driving expression in all photoreceptors. 5’ and 3’ homology arms of around 1 Kb upstream and downstream of sgRNA1 and sgRNA2, respectively were cloned into this vector for homologous recombination. Both plasmids were injected into *y^1^ M{nos-Cas9.P}ZH-2A w\** embryos (BDSC # 54591). F1 progeny was screened for *mCherry* expression using a fluorescent stereomicroscope, and positive flies were confirmed by PCR (for the absence of *mmt* ORF and against non-specific insertion of the knock-in cassette) and genome sequencing of the whole locus. We also collected flies without an insertion event to be used as background controls. PCR and sequencing also confirmed these. The obtained stocks were crossed to *w*, hs-cre; Sco/CyO; Dr/TM3* flies to remove the *mCherry* cassette flanked by loxP sites. To generate a line expressing *Gal4* from the *mmt* locus (Suppl. Fig 5), we took advantage of the attP site present in the *mmt* knock-out flies and used an RMCE-based strategy to recombine in a vector via its attB site. This vector contains a *Gal4* sequence and the upstream and downstream sequences to restore the 5’ and 3’ regions that were removed in the *mmt* knock-out. A *mini-white* gene was also included in the vector as a selective marker. F1 progeny was screened for red eye color, and positive flies were confirmed by PCR and sequencing. We also collected flies without an insertion event to be used as background controls. PCR and sequencing also confirmed these. The obtained stocks were crossed to *w*, hs-cre; Sco/CyO; Dr/TM3* to remove the *mini-white* cassette flanked by loxP sites.

### Generation of *UAS*-*mmt*, *UAS*-*hNPB* and *UAS*-*hNPW* lines

*UAS*-*mmt* stocks were generated by amplifying the *mmt* ORF from the corresponding cDNA and cloning it into pUASg.attB using the Gateway System (Invitrogen) and injected into embryos of *yw, M(eGFP, vas-int, dmRFP)ZH-2A; P{CaryP}attP40* flies. Flies without successful insertion of the UAS cassette were used to generate stocks for background controls. DNA encoding the Human isoforms of *neuropeptide B* (based on the sequences NP-202, NM_148896.5) or *neuropeptide W* (based on the sequences NPW-201, NM_001099456.3) were chemically synthesized (GenScript) and *UAS*-*NPB* and *UAS*-*NPW* were generated using the same protocol described for *UAS*-*mmt*.

### flyPAD assays

flyPAD assays were performed as described in (Itskov et al., 2014). For food choice experiments, single flies in different dietary conditions were tested in arenas containing two types of food patches: 10% yeast or 20 mM sucrose, each mixed with 1% agarose.

For all experiments, flies were individually transferred to fly-PAD arenas by mouth aspiration and allowed to feed for 1 h at 25°C (or 30°C for TrpA1 experiments), 70% relative humidity. The total number of sips per animal was calculated using previously described flyPAD algorithms (Itskov et al., 2014). Non-eating flies (defined as having fewer than two activity bouts during the assay) were excluded from the analysis.

### Total mRNA extraction, RT-PCR, and quantitative real– time PCR

Flies used for mRNA extraction were snap-frozen in dry ice and kept at −80°C until head processing. Behavioral assays were performed in parallel to confirm that sibling flies presented the expected feeding phenotype. Fly heads were separated from other body parts by vortexing the Eppendorf tubes and posteriorly passing the debris through 710-mm and 425-mm sieves (Retsch GmbH). Fly heads were counted before homogenization to ensure that the same number was used for all replicates.

mRNA was extracted from heads of flies (three replicates per condition) using the following procedure: samples were ground and homogenized for 20 s (using pestles #Z359947, Sigma) in 100 μl of PureZOL (#732±6890, Bio-Rad). 250 μl of PureZOL was further added and mixed by pipetting and incubated at RT for 10 min. Finally, 350 μl of 100% ethanol was added, and the samples were mixed and transferred to a Zymo column (Direct-zol RNA MicroPrep #R2062, Zymo research). The manufacturer’s instructions were followed to purify the mRNA (including DNAse treatment), and samples were eluted in 50 μl of distilled RNase/DNase-free water.

The concentration of the total mRNA samples was determined by performing a spectrophotometer scan in the UV region. Total RNA (1 μg) was reverse transcribed (RT) using the iScript Reverse Transcription Supermix for RT-PCR kit (#170±8840 Bio-Rad), following the manufacturer’s instructions. Each cDNA sample was amplified using SsoFast EvaGreen Supermix on the CFX96 Real-Time System (Bio-Rad). Briefly, the reaction conditions consisted of 1 μl of cDNA, 1 μl (10 μM) of each primer, 10 μl of supermix, and 7 μl of water. The cycle program consisted of enzyme activation at 95°C for 30 s, 39 cycles of denaturation at 95°C for 2 s, and annealing and extension for 5 s. The primers used in this reaction are listed in Table 2. This experiment was performed using three experimental replicates and two technical replicates per genotype. Appropriate non-template controls were included in each 96-well PCR reaction, and dissociation analysis was performed at the end of each run to confirm the specificity of the reaction. Absolute RNA levels were calculated from a standard curve and normalized to the internal controls (*Actin42A* and *RpL32*). Data processing was performed using Bio-rad CFX Manager 3.1 (Bio-Rad).

### Immunohistochemistry, image acquisition, and analysis

For visualization of expression patterns, males from each *Gal4* line were crossed to females homozygous for the *UAS-CD8∷GFP* reporter. Mated female flies with both the *Gal4* and the UAS reporter were kept on a complete holidic medium for 3 days prior to dissections. Samples were dissected in 4°C PBS, then transferred to formaldehyde solution (4% paraformaldehyde in PBS + 10% Triton-X) and incubated for 20–30 min at room temperature. Samples were washed three times in PBST (0.5% Triton-X in PBS) and then blocked in normal goat serum 10% in PBST for 60 min at room temperature. Primary antibodies used were: Rabbit anti-GFP (TP401, Torrey Pines Biolabs) at 1:6000; Mouse anti-GFP (A11120, Invitrogen) 1:500; Rabbit anti-Mmt (Eurogentec, generated in this study, see below); and Mouse anti-NC82 (nc82 was deposited to the DSHB by Buchner, E., Developmental Studies Hybridoma Bank) at 1:10. All primary antibodies were prepared in 5% normal goat serum in PBST and incubations were performed for 3 days at 4°C with shaking. They were then washed in PBST 2–3 times for 10–15 min at room temperature. The secondary antibodies used were Anti-Mouse 488 (A11029, Invitrogen), Anti-mouse A594 (A11032, Invitrogen), Anti-rabbit A488 (A11008, Invitrogen), Anti-Rabbit A594 (A11037, Invitrogen) at 1:500 in 5% normal goat serum in PBST and samples were then incubated for 3 days at 4°C with shaking. They were again washed in PBST 2–3 times for 10–15 min at room temperature. Samples were mounted in Vectashield (H-1000, Vector Laboratories). Images were captured on a Zeiss LSM 980 confocal microscope using a 20x or 40x objective, and Fiji (Schindelin et al., 2012) was used for imaging analysis.

### Generation of Mmt Antibody

To visualize the localization of Mmt in the nervous system, a custom anti-rabbit polyclonal antibody was generated by the company Eurogentec as a service. In brief, a peptide of 15 amino acids (EVEQD-VAGDQSAGRS) between E55 and S69 of the Mmt amino acid sequence was synthesized to be used as an antigen. The service company followed an 87-day immunization protocol and the final serum was purified using an affinity column. Upon optimization, we chose a dilution of 1:1000 to be used as the concentration for histochemistry.

### Video-tracking assay

Video tracking foraging assays were performed based on previously described protocols (Corrales-Carvajal et al., 2016; Münch et al., 2022). In brief, flies were individually transferred to circular arenas by mouth aspiration and allowed to freely forage for one hour at 25°C, 70% RH, while being video-recorded. Each arena was prepared with 12 equally distributed food spots, six containing 100 mM sucrose and six 10% yeast, each in 1% agarose. The Bonsai framework was used to acquire the data (Lopes et al., 2015), and analysis was done in Python. Based on the validation in Corrales-Carvajal et al. (2016), we used micromovements as a proxy for feeding behavior.

### Sequence and phylogenetic analysis

Relaxed (Expect threshold = 100) BLASTp searches against all non-redundant GenBank CDS translations+PDB+SwissProt+PIR+PRF excluding environmental samples from WGS projects were used to find proteins with similarities to CG14075 (NCBI Reference Sequence: NP_649080.1). This search yielded hits only from relatively closely-related brachyceran flies, which were aligned using MUltiple Sequence Comparison by Log-Expectation, MUSCLE (Edgar, 2004) (https://www.ebi.ac.uk/Tools/msa/). This alignment was used to identify conserved features in brachyceran CG14075 (Mmt) sequences and to derive the relaxed motif M-X(20,80)-W-{C}(20,80)-C-{CH}(9,13)-C-{CH}(9,13)-C-{CH}(9,13)-C. Signal peptides were predicted using SignalP 6.0 (https://dtu.biolib.com/SignalP-6) (Teufel et al., 2022). The general motif (H*)R/K-Xn-R/K, where X is any amino acid, n is 0, 2, 4, or 6 amino acids, and an H* denotes a histidine is accepted when n is 2, 4, or 6, was used to annotate putative protein cleavage sites (Devi, 1991; Hook et al., 2008; Seidah & Chrétien, 1999; Veenstra, 2000).

To identify Mmt orthologues outside of brachycera, a pattern search against Eukaryote KEGG genes was performed using the Motif search tool from the Kyoto University Bioinformatics Center (https://www.genome.jp/tools/motiftools/motif/MOTIF2.html). Hits were retained if they had the motif mentioned above, had a signal peptide, were less than 200 AA in length, and were not an obvious member of an annotated protein family with superior reciprocal blast hits in *Drosophila melanogaster*. The sole protein satisfying these rules for *Homo sapiens* was Neuropeptide B (NP_683694.1), which was not considered to have orthologues outside of Deuterostomia. After this identification, relaxed BLASTp (or TBLASTN) searches were run with CG14075 or NPB propeptides as queries to identify further hits, following the same selective criteria described above. Divergent hits from distantly-related taxa were selected as queries, and relaxed-BLASTp (or TBLASTN) searches were exhaustively repeated until no new clade could be identified. Considering all identified Mmt-like sequences, the consensus motif for the Mmt-like proteins was refined to <M-X(15,30)-{C}(4,110)-C-{CW}(2)-[DE]-{W}(7,12)-C-{C}(8,12)-C-{C}(11,14)-C-{C}(1,50)>. The Marmite domain is defined as the C-terminal portion of Marmite-like proteins containing the 4 cysteines.

For phylogenetic analysis, a tree was created with the Simple phylogeny tool available (https://www.ebi.ac.uk/Tools/phylogeny/simple_phylogeny/) using CG14075-like, NPB-like and NPW-like prepropeptide sequences aligned using the T-COFFEE multiple sequence alignment program with the BLOSUM62 matrix (https://www.ebi.ac.uk/Tools/msa/) (Notredame et al., 2000). CBP family proteins were included as outgroups. Sequences were then clustered using the neighbor-joining algorithm to construct trees from the distance matrix with gap included and with distance correction to correct for multiple substitutions at the same site (Kimura, 1980; Saitou & Nei, 1987). Unrooted trees were displayed, annotated, and managed using The Interactive Tree Of Life tool (https://itol.embl.de/) (Letunic & Bork, 2021).

Sequence alignments were visualized and edited using Jalview (Waterhouse et al., 2009). All sequences used for the phylogenetic analysis are listed in Table 3.

### Mice

All experiments were approved by the Champalimaud Foundation and Portuguese Direção Geral de Veterinária, and performed in accordance with the European Union Directive for Protection of Vertebrates Used for Experimental and other Scientific Ends (86/609/CEE). Male and female C57BL/6J mice were used from the inbred colony of the Champalimaud Foundation Vivarium. All animals started the procedure at 8-9 weeks of age.

### Surgical procedures and viral Injections in mice brain

C57BL/6J mice were anesthetized using a mix of oxygen (1-1.5 L/min) and 1-3% isoflurane (Vetflurane, Virbac). Briefly, the hair on the top of the head was clipped, the mouse was placed on a surgical table, covered with a heating pad set at 37°C and stabilized in the stereotaxic apparatus (David Koft Instruments). A skin incision was performed to expose the skull, connective and muscle tissue was carefully removed, and the skull surface was leveled at less than 0.05 mm by comparing the height of bregma and lambda, and medial-lateral directions. Two small holes were drilled for bilateral viral injections. Bilateral viral injections were performed using glass pipettes with 3μl *marmite (mmt)*, *neuropeptide B* (*NPB*) or control viral stock solution. *mmt*: AAV5-hSyn-*dmmt*-2A-*mCherry*; *NPB*: AAV5-hSyn-*mNPB*-2A-*mCherry*; *Ctrl*: AAV.CAG.Flex.GCaMP6f.WPRE.SV40 (AAV5) (Addgene 100835) or AAVsyn1-tdTomato-WPRE(AAV5) (Addgene 51506). *mmt* and *NPB-*expressing viruses were produced by the Molecular and Transgenic Tools Platform at Champalimaud Foundation.

0.5 μl of virus were bilaterally injected in the Ventromedial Hypothalamus (VMH) (Antero-posterior (AP)-1.3mm; Medio-lateral (ML): +-0.3mm and Dorso-ventral (DV):-5.6 mm (from brain surface; Paxinos 2008), at a rate of 0.96 nL/s using nanoject II (Drummond Scientific), with the pipette retracted only twenty minutes after the injection was completed. After the surgery, animals recovered in their home cage with a heating pad set at 37°C and were closely monitored until they fully recovered.

### Food intake measurements of mice

C57BL/6J animals were first separated one per cage and allowed an acclimatization period of 1 week to ensure weight stabilization. Food intake measurements were performed every 2 days over 10 weeks: 2 weeks before surgery (baseline measurements) and 8 weeks after *mmt*, *NPB* or Ctrl viral injections. 2014 Food pellets of Teklad global 14% protein rodent maintenance diets (Envigo) were used. Food intake was deduced by measuring the difference between the initial and final weight of the provided food using a scale (Ohaus bench scale).

### Histology and immunohistochemistry of mice samples

After feeding measurement experiments were completed, mice were anesthetized with an intraperitoneal injection of ketamine-xylazine (100 mg/kg and 5 mg/kg, respectively), and perfusion was performed with 1x phosphate buffered saline (PBS) and 4 % paraformaldehyde (PFA). Brains were gently extracted and placed in 4% PFA overnight and then transferred to PBS at 4°C for further histological processing. Brains were sectioned coronally in 50 μM slices using a vibratome (Leica VT1000S), and sections were collected into a 24-well plate with PBS before mounting. After mounting, images of the sections were taken using an automated wide-field fluorescence microscope (Zeiss AxioScan.Z1). The acquisition features were standardized for all the slices and acquired with Plan - Apochromat 10 X/0.45 M27 (Zeiss). A green and a red wavelength laser (channel Alexa Fluor (AF) 488, channel red (Cy3) 570, excitation filter 555 nm) were used with exposure times of 50 ms and 75ms, respectively, and a LED power of 10 %. Finally, the output images were transformed into 2D maximum intensity images using the ZEN 3.4 edition (Zeiss) software.

### Statistical analysis

All violin plots show the frequency distribution of the data, with lines representing the median, upper and lower quartile. In all behavioral experiments, each data point corresponds to one animal, and the sample sizes correspond to the number of animals tested. All statistical tests used were two-sided. Normality was tested using the Shapiro-Wilk normality test, and parametric or nonparametric tests were used accordingly. The statistical tests used in each experiment are stated in the figure legends.

## Author Contributions

CR and APF conceived and designed the project, performed data analysis, interpreted data, and wrote the original draft of the manuscript; APF, IT and ABF performed experiments and data analysis; GE-N, BD, AJO-M and AMG performed data analysis; CR, BD, AJO-M and AMG contributed through project supervision, administration, and funding acquisition; all authors reviewed and edited the manuscript.

