## Supplementary Information for "*marmite* defines a novel conserved neuropeptide family mediating nutritional homeostasis"

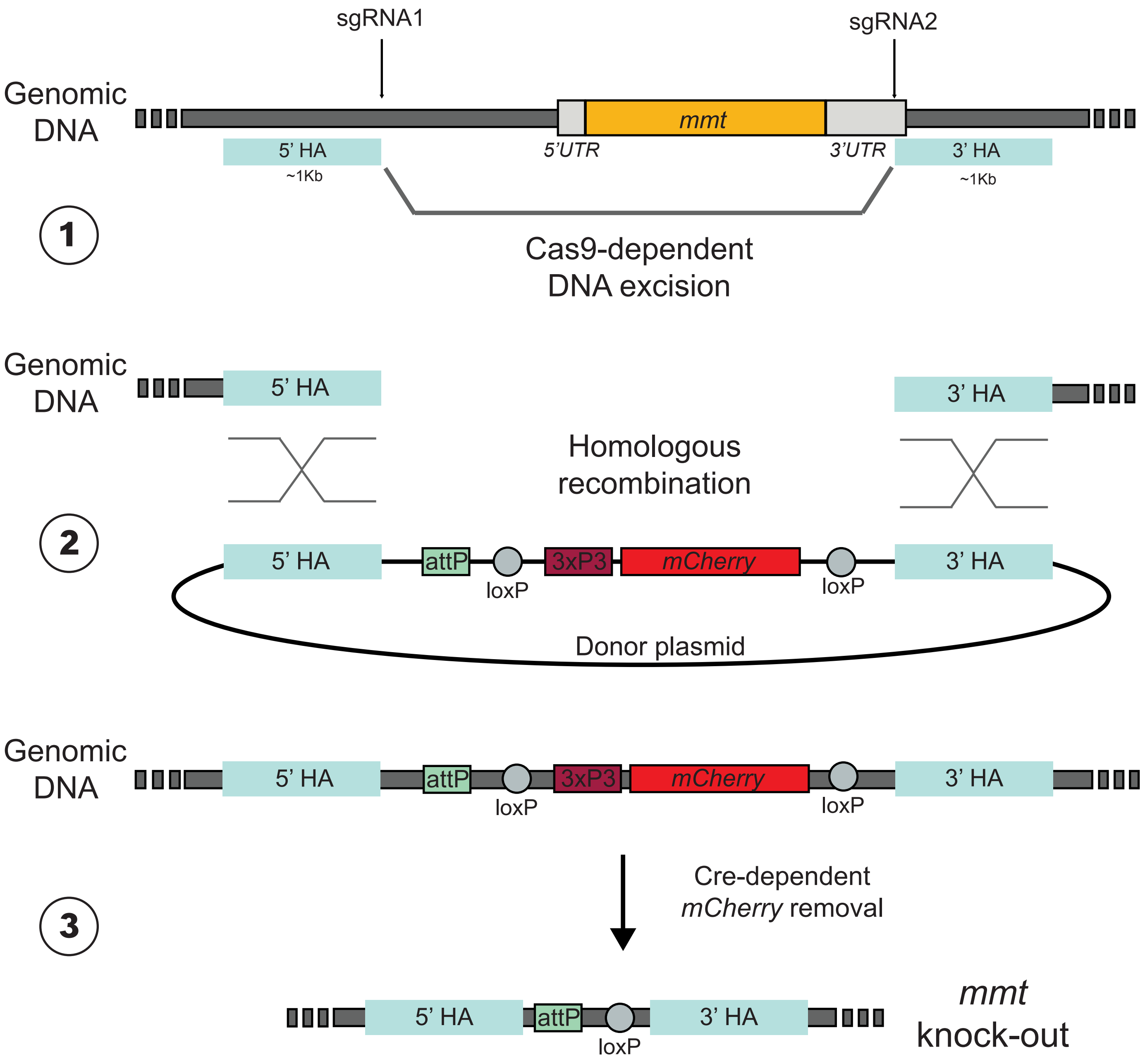


**Supplementary Figure 1 – Generation of *mmt* knock-out.** Scheme depicting the generation of the *mmt* knock-out line using CRISPR-Cas9 genome editing techniques. **Step 1:** Cas-9-dependent DNA excision between the 5' and the 3' homology arms (HA) mediated by two single guide RNAs (sgRNA). **Step 2:** A donor plasmid, including an attP site for later insertions of transgenes and a *mCherry* as a selective marker under the 3xP3 promoter driving expression in all photoreceptors, was inserted by homologous recombination into the excised part of the genome between the 5' and 3' homology arms (HA). **Step 3:** The *mCherry* cassette was removed by inducing a Cre-mediated recombination event between the loxP sites flanking the cassette.

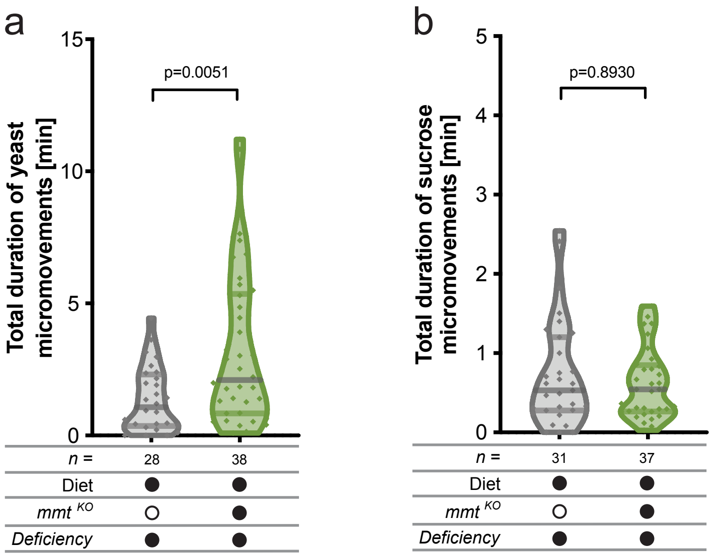


**Supplementary Figure 2 – *mmt* mutant flies increase the total duration of yeast but not sucrose micromovements.** Total duration of yeast **(a)** and sucrose **(b)** micromovements of *mmt* mutants and control flies. “Deficiency” denotes a genetic deficiency spanning the *mmt* gene. Flies were maintained on complete holidic medium. Below the graphs, filled black circles represent the complete holidic medium or presence of a specific genetic element, whereas an empty black circle represents its absence. Violin plots show the frequency distribution of the data, with lines representing the median, upper and lower quartiles; points indicate the total duration of micromovements on yeast **(a)** or sucrose **(b)** for each fly. Significance was tested using the Mann-Whitney test. The number of flies used in each condition is shown in each panel.

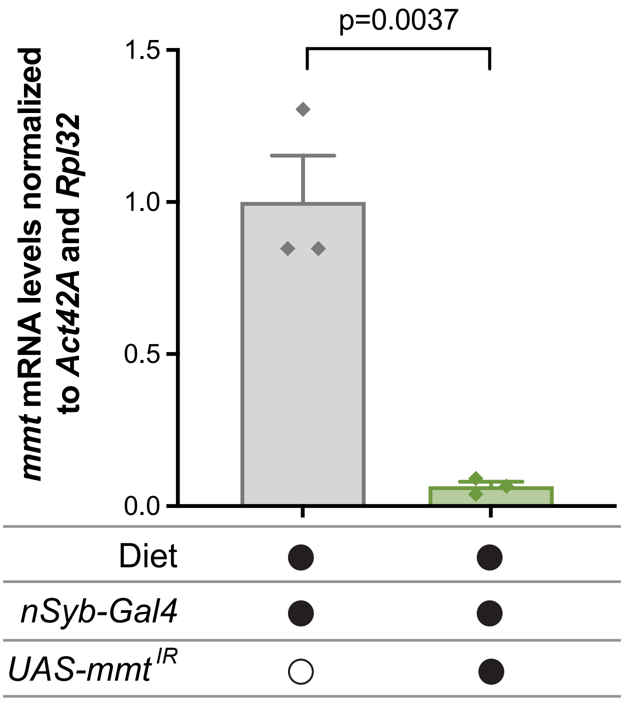


**Supplementary Figure 3 – Neuronal knockdown of *mmt*.** *mmt* mRNA levels measured from heads of *mmt* pan-neuronal knockdown or control flies maintained on complete holidic medium as normalized using the expression levels of two control genes (*Actin 42A* and *RpL32*). Filled black circles represent the complete holidic medium or presence of a specific genetic element, whereas an empty black circle represents its absence. The columns represent the mean and the error bars the standard error of the mean. Significance was tested using the unpaired t-test, n=3. Related to Figure 1d.

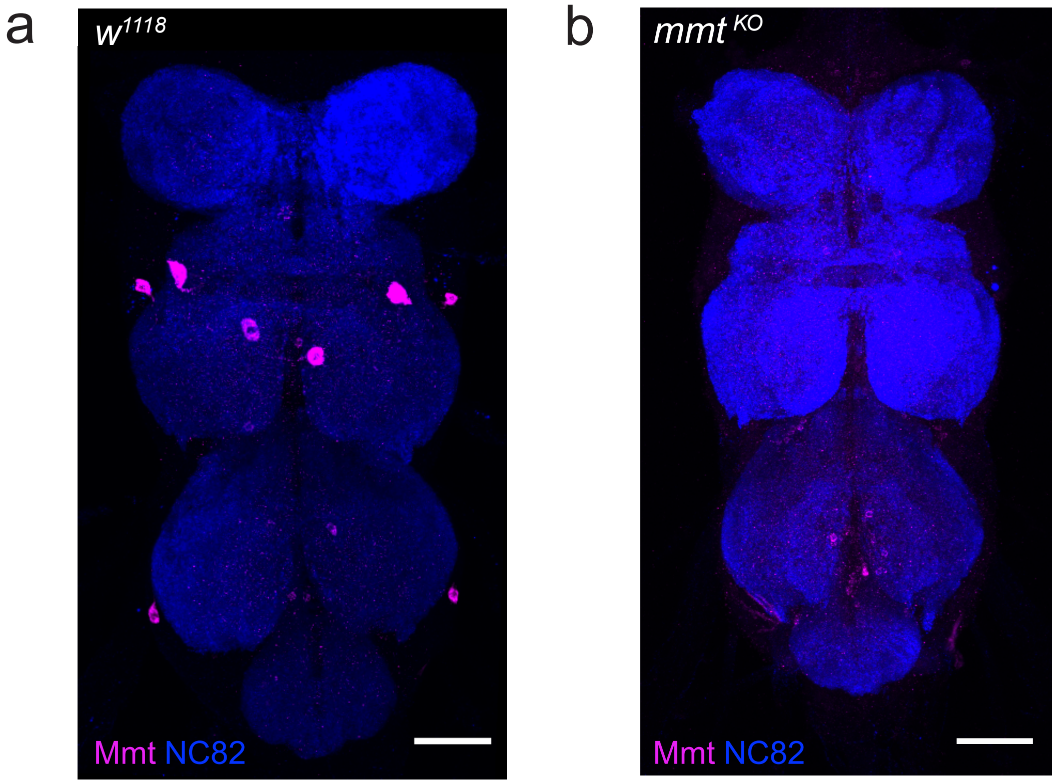


**Supplementary Figure 4 – Mmt antibody is expressed in the ventral nerve cord.** Mmt antibody staining of VNC of control **(a)** and *mmt* mutant **(b)** flies in magenta, with nc82 neuropile staining in blue. VNCs are oriented with the anterior side on top and the posterior side at the bottom. Maximum projection images of confocal stacks; scale bars correspond to 50 µm.

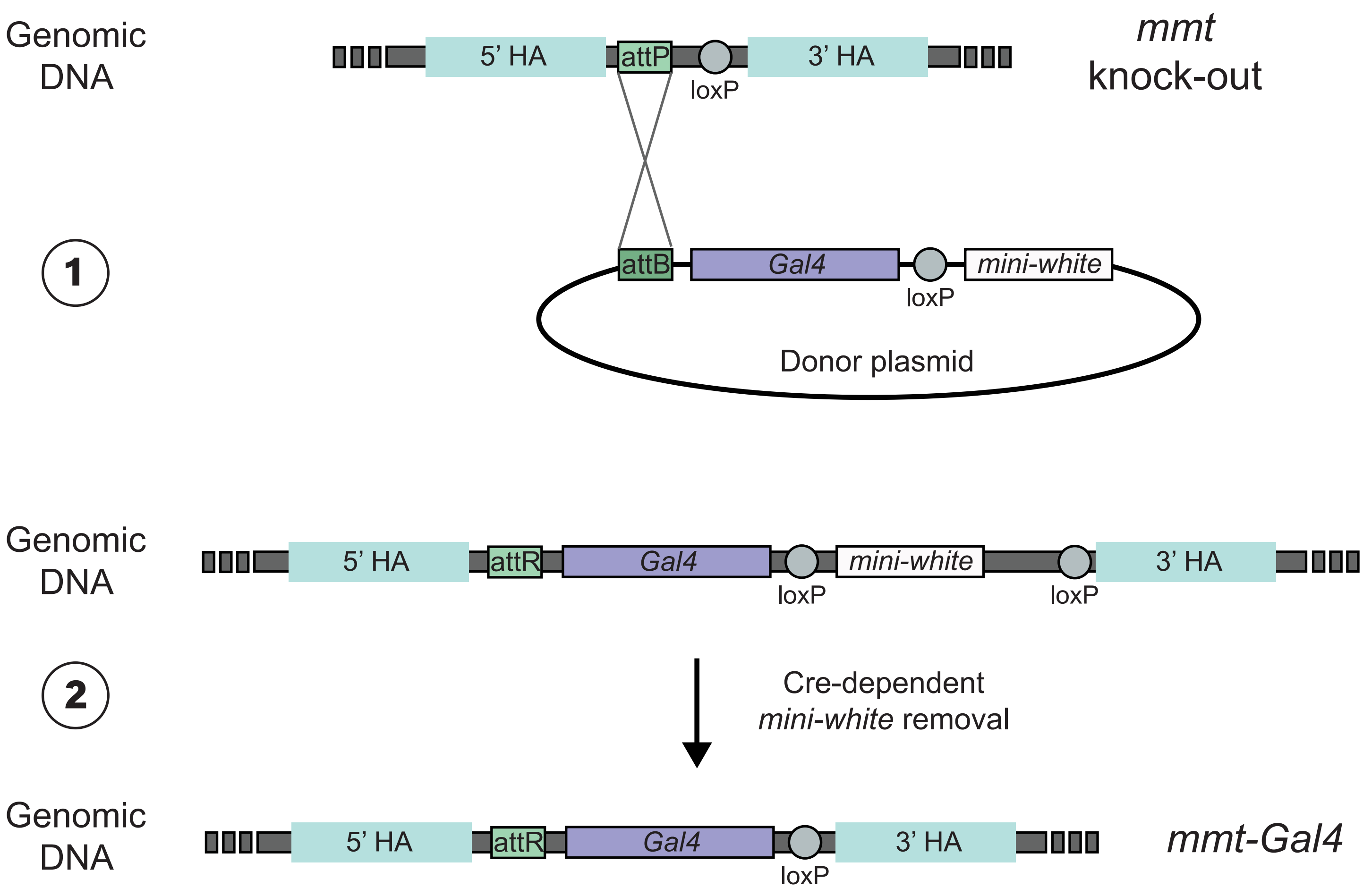


**Supplementary Figure 5 – Generation of mmt-Gal4 knock-in.** Scheme depicting the generation of *mmt-Gal4* knock-in. **Step 1:** A donor plasmid carrying a *Gal4* and a *mini-white* as a selection marker was inserted into the *mmt* knock-out locus via ΦC31-recombinase-mediated cassette exchange. **Step 2:** The *mini-white* cassette was removed by inducing a Cre-mediated recombination event between the loxP sites flanking the cassette.

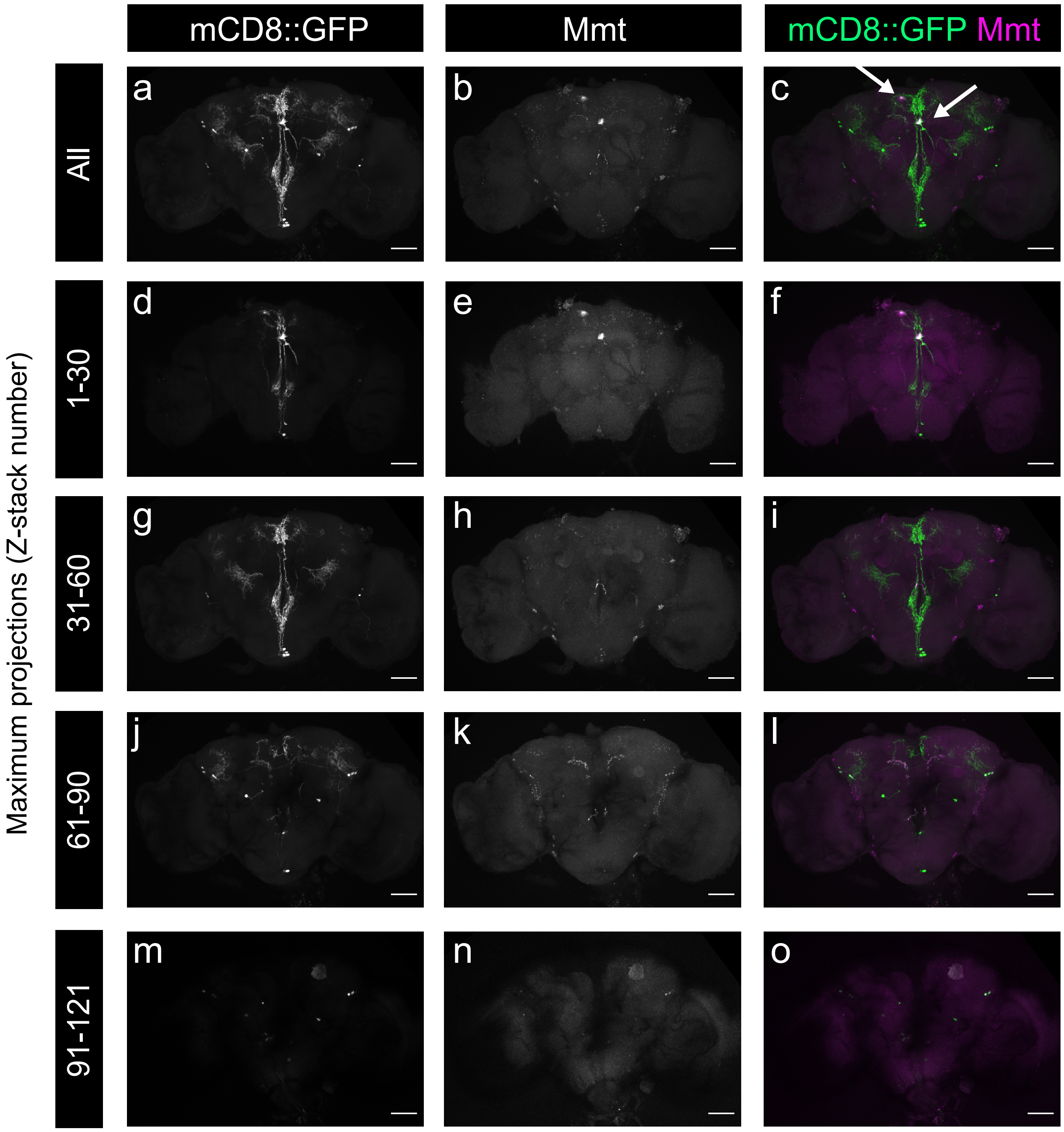


**Supplementary Figure 6 – *mmt-Gal4* recapitulates Mmt antibody expression. a-o)** Maximum projections of all or sub-sections of confocal stacks of brains of *mmt-Gal4* flies driving *UAS-mCD8::GFP*, showing mCD8::GFP, Mmt antibody or both; scale bar corresponds to 50 μm.

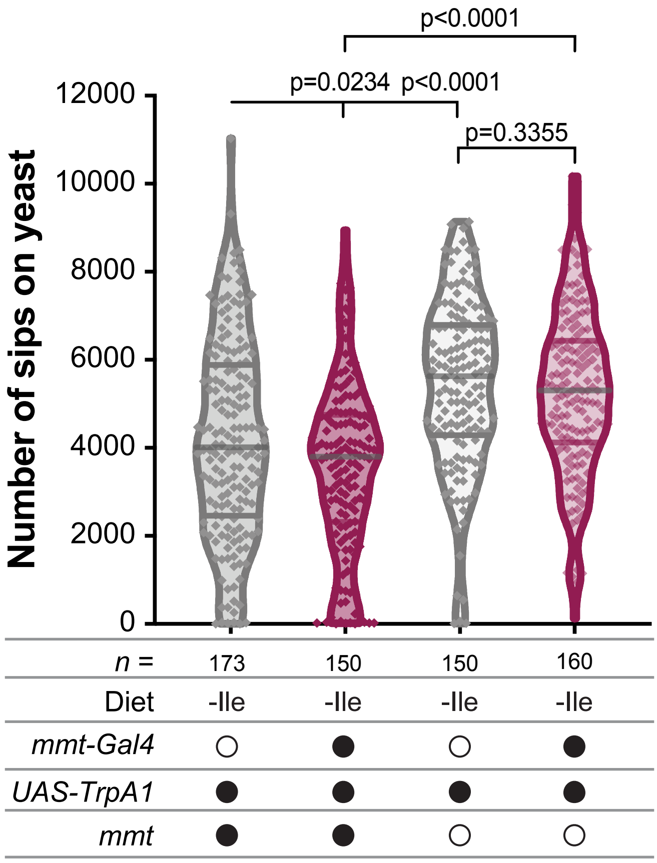


**Supplementary Figure 7 – Activation of *mmt-Gal4* expressing neurons.** Full dataset of the number of sips on yeast in which *mmt-Gal4* neurons were activated in a context of *mmt* presence or absence and corresponding controls. Flies were maintained on holidic medium lacking Isoleucine (-Ile). Related to Figure 3b. Below the graph, filled black circles represent the presence of a specific genetic element, whereas an empty black circle represents its absence. Violin plots show the frequency distribution of the data, with lines representing the median, upper and lower quartiles; points indicate the number of sips on yeast for each fly. Significance was tested using the Mann-Whitney test. The number of flies used in each condition is shown in each panel.

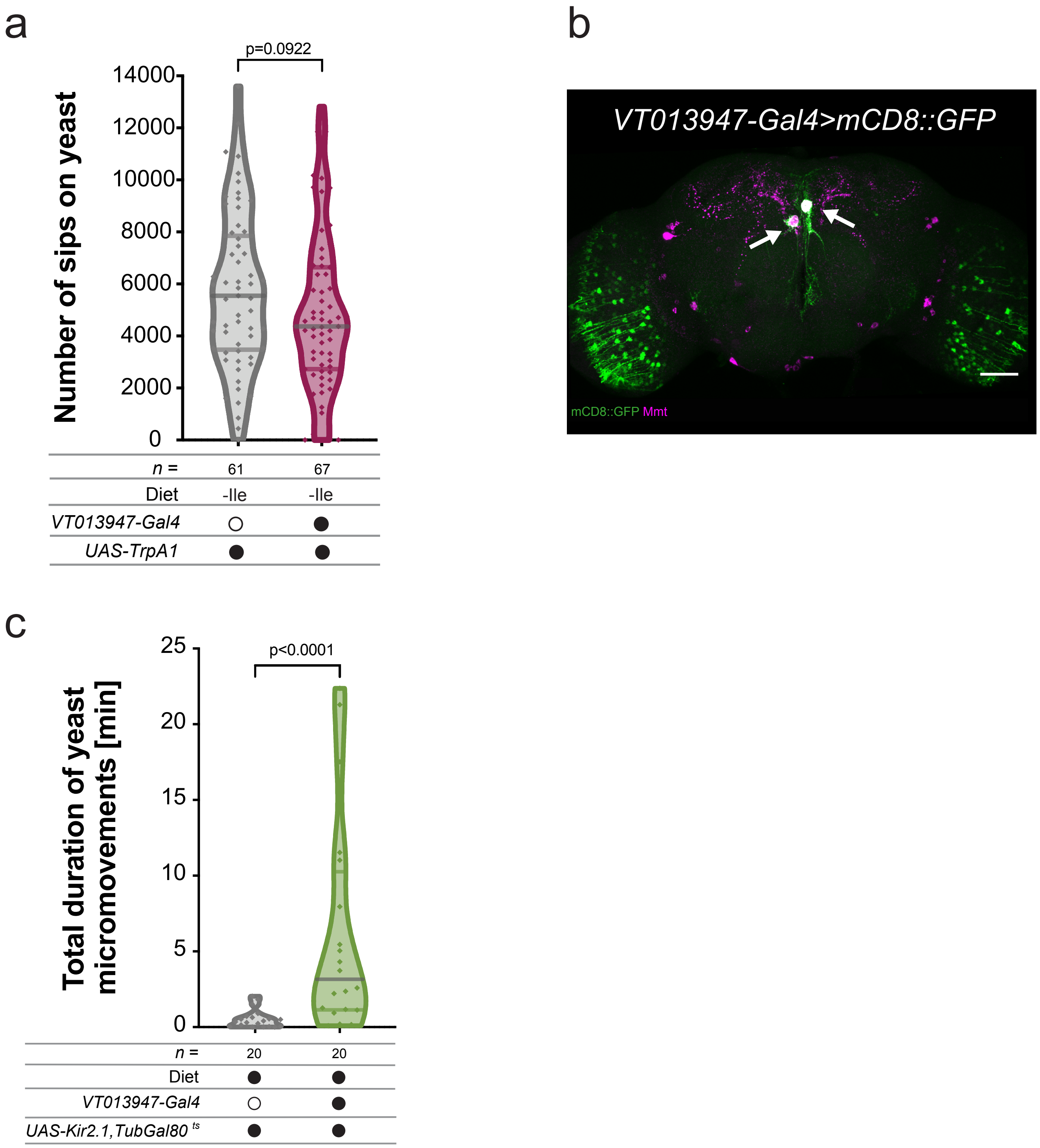


**Supplementary Figure 8 – *VT013947-Gal4* neurons modulate protein appetite. a)** Full dataset of the number of sips on yeast of flies in which *VT013947-Gal4* neurons were activated and corresponding control. Flies were maintained on holidic medium lacking Isoleucine (-Ile). Related to Figure 3d. **b)** Expression pattern of *VT013947-Gal4* in the brain visualized using *UAS-mCD8::GFP* in green, co-stained with Mmt antibody in magenta. The brain is oriented with the dorsal side on top and the ventral side at the bottom. Maximum projection images of confocal stacks; scale bar corresponds to 50 µm. **c)** Total duration of yeast micromovements of flies in which *VT013947-Gal4* neurons were inducibly silenced using *Kir2.1* and corresponding control. Flies were maintained on complete holidic medium. Below the graphs, filled black circles represent the complete holidic medium or presence of a specific genetic element, whereas an empty black circle represents its absence. Violin plots show the frequency distribution of the data, with lines representing the median, upper and lower quartiles; points indicate the number of sips on yeast **(a)** or total duration of yeast micromovements **(c)** for each fly. Significance was tested using the Mann-Whitney test. The number of flies used in each condition is shown in each panel.

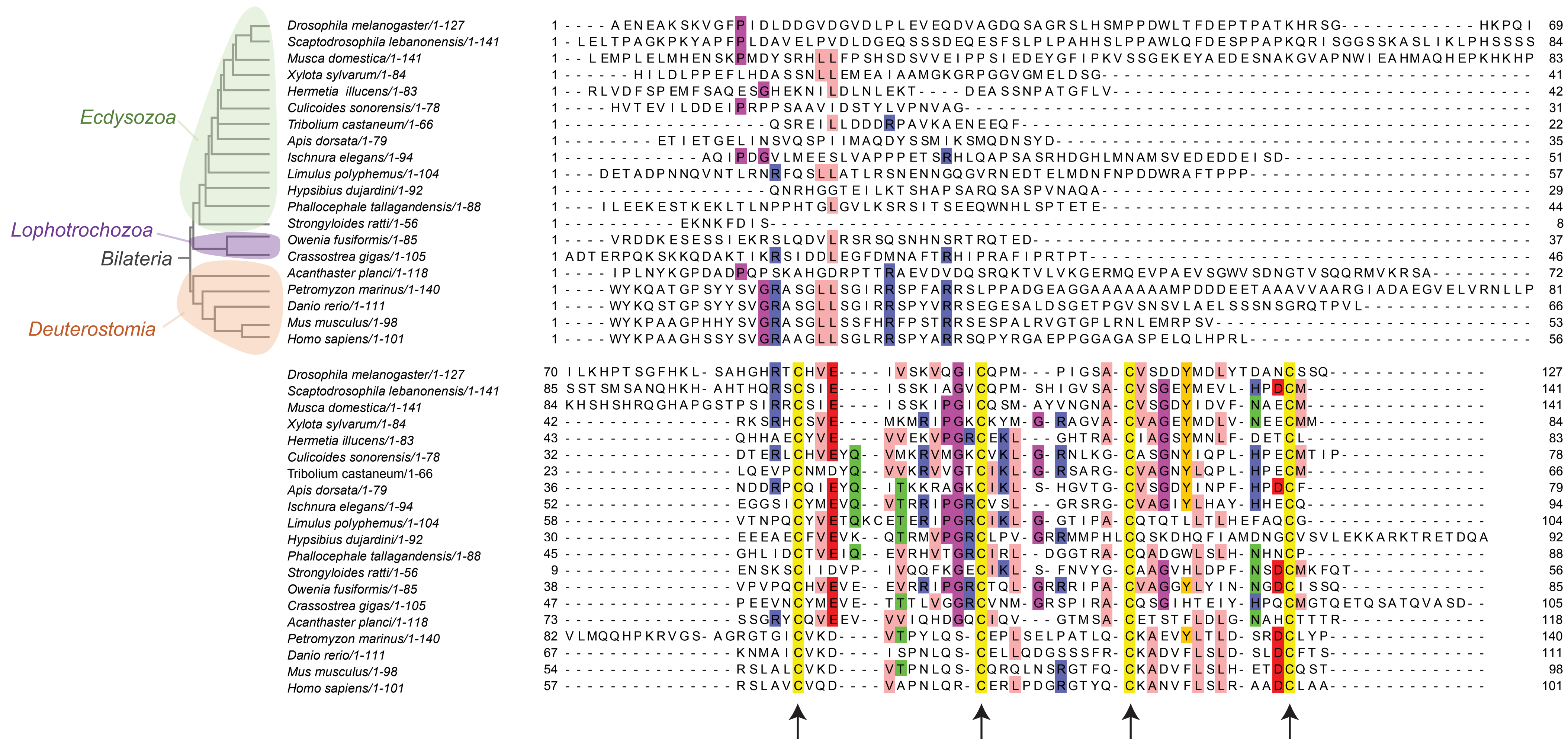


**Supplementary Figure 9 – Alignment of Mmt-related and NPB AA sequences without signal peptide.** Alignment of Mmt-like and NPB AA sequences without the predicted signal peptides from invertebrate and vertebrate species. The main taxa to which the different species belong are shown. Each of the four conserved cysteines in the C-terminal part is highlighted with an arrow. Alignment representing 30% identity.

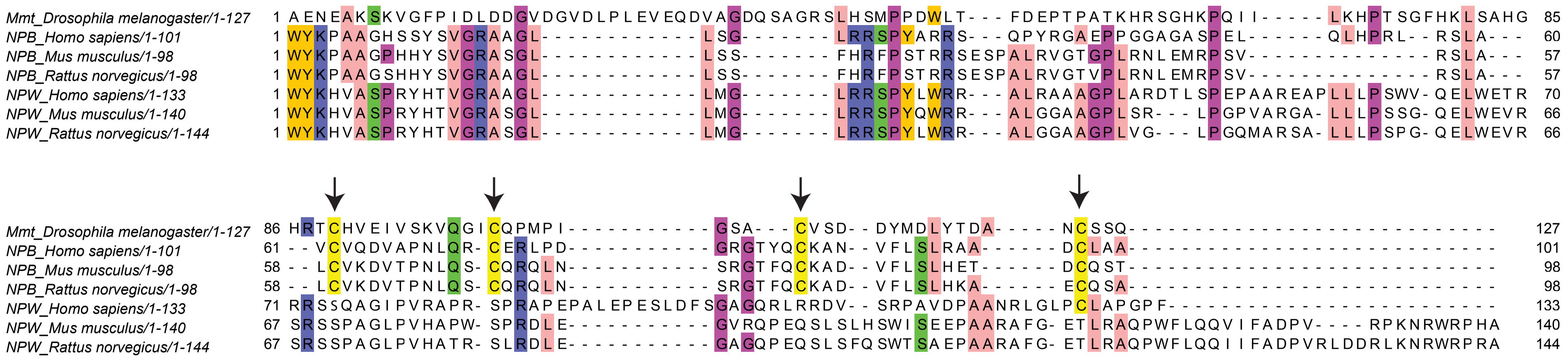


**Supplementary Figure 10 – Alignment of Mmt, NPB and NPW AA sequences without signal peptide.** Alignment of *Drosophila* Mmt, and NPB/W AA sequences from human, mouse and rat without the predicted signal peptides. Each of the four conserved cysteines in the C-terminal part is highlighted with an arrow. Alignment representing 50% identity.

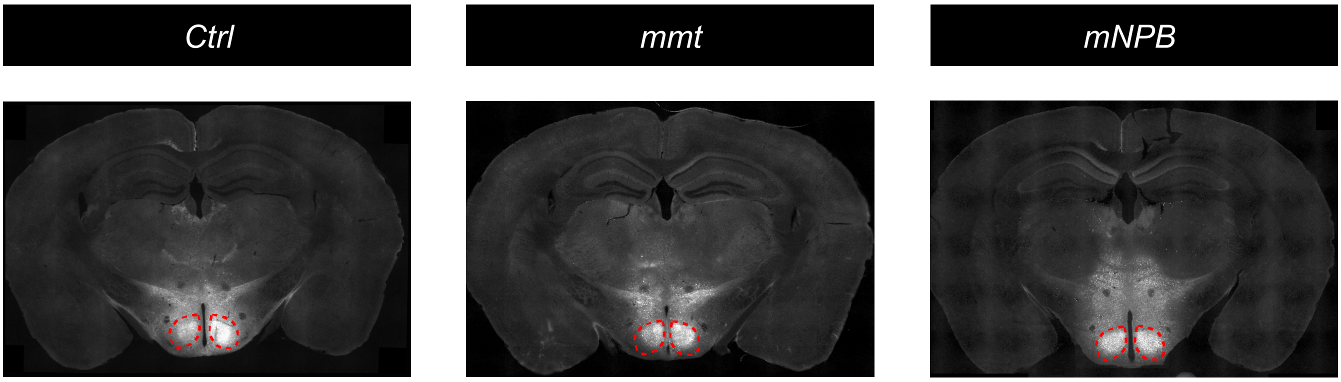


**Supplementary Figure 11 – mmt and NPB-virus expression in mice.** Representative image of brains of mice injected with control (left), *mmt*-expressing virus (middle) or *NPB*-expressing virus (right) stained for fluorescent proteins. A dashed red line indicates the site of injection (VMH).

### Table 1 - List of stocks used in this study.

| Line | Full genotype | Source |
|---|---|---|
| *$w^{1118}$* | *$w^{1118}$; +; +* | Ribeiro Lab |
| *nos-Cas9* | *$y^1$ M{nos-Cas9.P}ZH-2A w** | BDSC #54591 |
| *hs-cre* | *w*,hs-cre; Sco/CyO; Dr/TM3* | Chiappe Lab |
| *mmt $^{KO}$* | *$w^{1118}$; +; $mmt^{KO\ 30.2.1}$* | this study |
| *background control for mmt $^{KO}$* | *$w^{1118}$; +; +* | this study |
| *Deficiency* | *$w^{1118}$;+; Df(3L)BSC220/TM6c* | BDSC #9697 |
| *nSyb-Gal4* | *$w^{1118}$ , UAS-DCR2/Yhs-hid; +; nSyb-Gal4 2.1/ TM6b* | Ribeiro Lab |
| *UAS-mmt $^{IR}$* | *$w^{1118}$, P{GD6177}v14328;* | VDRC #14328 |
| *nSyb-Gal4* | *$w^{1118}$/Yhs-hid; +; nSyb-Gal4 2.1/ TM6b* | Ribeiro Lab |
| *attP40* | *y w M(eGFP, vas-int, dmRFP)ZH-2A; P{CaryP}attP40* | Champalimaud Fly platform |
| *background control for UAS-mmt, hNPB and hNPW* | *$w^{1118}$; +;+* | this study |
| *UAS-mmt* | *$w^{1118}$; UAS-$mmt^{8.1}$/CyO* | this study |
| *UAS-hNPB* | *$w^{1118}$; UAS-$hNPB^1$/CyO* | this study |
| *UAS-hNPW* | *$w^{1118}$; UAS-$hNPW^1$/CyO* | this study |
| *mmt-Gal4* | *$w^{1118}$; +; mmt $^{Gal4\ 13.1.1}$ /TM6b* | this study |
| *background control for mmt-Gal4* | *$w^{1118}$; +; mmt $^{KO\ 13.4.1}$ /TM6b* | this study |
| *UAS - mCD8::GFP* | *$w^{1118}$/Yhs-hid; mCD8::GFP* | Ribeiro Lab |
| *UAS - mCD8::GFP; UAS-redstinger* | *w*; mCD8::GFP; UAS-redstinger* | Moita Lab |
| *UAS-TrpA1* | *$w^{1118}$/Yhs-hid; UAS-TrpA1* | Ribeiro Lab |
| *UAS-TrpA1; Deficiency* | *$w^{1118}$; UAS-TrpA1/CyO; Df(3L)BSC220/TM6b* | this study |
| *UAS-kir2.1, $tubGal80^{ts}$* | *$w^*$/Yhs-hid ; + ; UAS-kir2.1::eGFP, $tubGal80^{ts}$/TM6b* | Vasconcelos Lab |
| *VT013947-Gal4* | *P{VT013947-Gal4}attP2* | VDRC #205650 |
| *empty-Gal4* | *$w^{1118}$; P{GAL4.1Uw}attP2* | BDSC#68384 |

### Table 2 - List of primers used for RT-qPCR.

| Primer | Sequence |
|---|---|
| *mmt Fwd* | 5' ATCTGCTGCTCCACTGTGC 3' |
| *mmt Rev* | 5' TTCGATTTGGCCTCATTTTC 3' |
| *Actin42A Fwd* | 5' CAGGCGGTGCTTTCTCTCTA 3' |
| *Actin42A Rev* | 5' AGCTGTAACCGCGCTCAGTA 3' |
| *RpL32 Fwd* | 5' GCCCAAGATCGTGAAGAAGC 3' |
| *RpL32 Rev* | 5' GCACTCTGTTGTCGATACCCTTG 3' |

**Table 3 - List of sequences used for phylogenetic analysis.**

| Reference | Species | Common name | Protein |
|---|---|---|---|
| AAF52668.1 | *Drosophila melanogaster* | Vinegar fly | CBP |
| NP_652325.2 | *Drosophila melanogaster* | Vinegar fly | CBP |
| KXJ76498.1 | *Aedes albopictus* | Asian tiger mosquito | CBP |
| XP_037906351.1 | *Hermetia illucens* | Black soldier fly | CBP |
| XP_018785622.1 | *Bactrocera latifrons* | Fruit fly | CBP |
| NP_001092926.2 | *Homo sapiens* | Human | NPW |
| XP_014350484.1 | *Latimeria chalumnae* | Coelacanth | NPW |
| NP_695206.1 | *Rattus norvegicus* | Rat | NPW |
| NP_001093134.1 | *Mus musculus* | Mouse | NPW |
| NP_683694.1 | *Homo sapiens* | Human | NPB |
| NP_001334545.1 | *Mus musculus* | Mouse | NPB |
| NP_001120841 | *Danio rerio* | Zebrafish | NPB |
| NP_695205.1 | *Rattus norvegicus* | Rat | NPB |
| XP_032826114.1 | *Petromyzon marinus* | Sea lamprey | NPB-like |
| NP_649080.1 | *Drosophila melanogaster* | Vinegar fly | CG14075/Mmt |
| XP_022103320.1 | *Acanthaster planci* | Crown-of-thorns starfish | Mmt-Like |
| CAC9654597.1 | *Owenia fusiformis* | Annelida | Mmt-Like |
| XP_024499497.1 | *Strongyloides ratti* | Nematode | Mmt-Like |
| TSA:_isolate_PT2011_Phta_Filter15_CLC_contig_27382 | *Phallocephale tallagandensis* | Onychophora | Mmt-Like |
| TSA:_transcribed_RNA_sequence_GBZR01002010.1 | *Hypsibius dujardini* | Tardigrade | Mmt-Like |
| XP_022243359.1 | *Limulus polyphemus* | Horseshoe crab | Mmt-Like |
| XP_046389499.1 | *Ischnura elegans* | Blue-tailed damselfly | Mmt-Like |
| XP_031368331.1 | *Apis dorsata* | Giant honey bee | Mmt-Like |
| KYB28121.1 | *Tribolium castaneum* | Red flour beetle | Mmt-Like |
| LN483579.1 | *Culicoides sonorensis* | Biting midge | Mmt-Like |
| CAD7080404.1 | *Hermetia illucens* | Black soldier fly | Mmt-Like |
| LR999959.1 | *Xylota sylvarum* | Hoverfly | Mmt-Like |
| XP_005177225.1 | *Musca domestica* | House fly | Mmt-Like |
| XP_030387299.1 | *Scaptodrosophila lebanonensis* | Fly | Mmt-Like |
| XP_019929219.2 | *Crassostrea gigas* | Giant Pacific Oyster | Mmt-Like |
